# The heptaprenyl diphosphate synthase (Coq1) is the target of a lipophilic bisphosphonate that protects mice against *Toxoplasma gondii* infection

**DOI:** 10.1101/2022.04.04.487054

**Authors:** Melissa A. Sleda, Zhu-Hong Li, Ranjan Behera, Baihetiya Baierna, Catherine Li, Jomkwan Jumpathong, Satish R. Malwal, Makoto Kawamukai, Eric Oldfield, Silvia N. J. Moreno

## Abstract

Prenyldiphosphate synthases catalyze the reaction of allylic diphosphates with one or more isopentenyl diphosphate molecules to form compounds such as farnesyl diphosphate, used in e.g. sterol biosynthesis and protein prenylation, as well as longer “polyprenyl” diphosphates, used in ubiquinone and menaquinone biosynthesis. Quinones play an essential role in electron transport and are associated with the inner mitochondrial membrane due to the presence of the polyprenyl group. In this work, we investigated the synthesis of the polyprenyl diphosphate that alkylates the ubiquinone ring precursor in *Toxoplasma gondii*, an opportunistic pathogen that causes serious disease in immunocompromised patients and the unborn fetus. The enzyme that catalyzes this early step of the ubiquinone synthesis is Coq1 (TgCoq1), and we show that it produces the C35 species, heptaprenyl diphosphate. TgCoq1 localizes to the mitochondrion, and is essential for *in vitro T. gondii* growth. We demonstrate that the growth defect of a *T. gondii* TgCoq1 mutant is rescued by complementation with a homologous *TgCoq1* gene or with a (C45) solanesyl diphosphate synthase from *Trypanosoma cruzi* (TcSPPS). We find that a lipophilic bisphosphonate (BPH-1218) inhibits *T. gondii* growth at low nM concentrations, while overexpression of the TgCoq1 enzyme dramatically reduced growth inhibition by the bisphosphonate. Both the severe growth defect of the mutant and the inhibition by BPH-1218 were rescued by supplementation with a long chain (C30) ubiquinone (UQ_6_). Importantly, BPH-1218 also protected mice against a lethal *T. gondii* infection. TgCoq1 thus represents a potential drug target that could be exploited for improved chemotherapy of toxoplasmosis.

**Importance:** Millions of people are infected with *Toxoplasma gondii* and the available treatment for toxoplasmosis is not ideal. Most of the drugs currently used are only effective for the acute infection and treatment can trigger serious side-effects requiring changes in the therapeutic approach. There is, therefore, a compelling need for safe and effective treatments for toxoplasmosis. In this work, we characterize an enzyme of the mitochondrion of *T. gondii* that can be inhibited by an isoprenoid pathway inhibitor. We present evidence that demonstrate that inhibition of the enzyme is linked to parasite death. In addition, the drug is able to protect mice against a lethal dose of *T. gondii*. Our results thus reveal a promising chemotherapeutic target for the development of new medicines for toxoplasmosis.

## Introduction

Apicomplexan parasites are responsible for important human and animal diseases including malaria and toxoplasmosis. The phylum member *Toxoplasma gondii* alone causes toxoplasmosis in aproximately one-third of the world’s population [1]. Most human infections are uncomplicated, but severe disease and death can occur in prenatal infections and in immunocompromised individuals [2]. In the United States there is an estimated 11% seroprevalence, with ∼1.1 million people infected each year with *T. gondii* [3]. Treatment for toxoplasmosis is challenged by the lack of an effective treatment for the chronic infection, and many patients do not respond to therapy [4]. Most of the drugs currently used are poorly distributed to the central nervous system and they can trigger serious side-effects requiring changes in the therapeutic approach [5]. There is, therefore, a compelling need for safe and effective treatments for toxoplasmosis.

The mitochondrion of *T. gondii* is essential for its survival and is a validated drug target as it houses important pathways like the Electron Transport Chain (ETC). Ubiquinone is an essential component of the ETC, and it is confined to the inner mitochondrial membrane by a long isoprenoid tail [6]. Ubiquinone biosynthesis in *T. gondii* is a relatively unexplored subject, but what it is known is that isoprenoid biosynthesis is essential for cell growth [7]. Isoprenoids are lipid compounds found in nature and they have many important functions [8, 9]. The enzymes that synthesize and use isoprenoids are among the most important drug targets for the treatment of cardiovascular disease, osteoporosis and bone metastases and have shown promise as antimicrobials [10–12]. The five carbon (C5) precursors of isoprenoids, isopentenyl diphosphate (IPP) and dimethylallyl diphosphate (DMAPP) (Fig. 1A) [13] are synthesized in mammalian cells by the mevalonate pathway, while other organisms—including apicomplexan parasites—use the 1-deoxy-D-xylulose-5-phosphate (DOXP) pathway [7](Fig. 1A, green square). The enzyme farnesyl diphosphate synthase (FPPS) synthesizes farnesyl diphosphate (FPP) from DMAPP and IPP and further elongation of FPP results in (C20) geranylgeranyl diphosphate (GGPP), a reaction catalyzed by GGPP synthases (GGPPS). Notably, *T. gondii* expresses a FPPS (TgFPPS) that synthesizes both FPP and GGPP [14]. We also previously showed that intracellular parasites are capable of acquiring GGPP from the host [15], making parasites resistant to bisphosphonate inhibitors of the *T. gondii* FPPS, such as zoledronate, but also sensitive to inhibition of the mammalian isoprenoid biosynthesis pathway [15]. It thus appeared possible that inhibiting the prenyldiphosphate synthase responsible for biosynthesis of the longer-chain species involved in ubiquinone biosynthesis might be an alternate therapeutic strategy.

**Figure 1.**
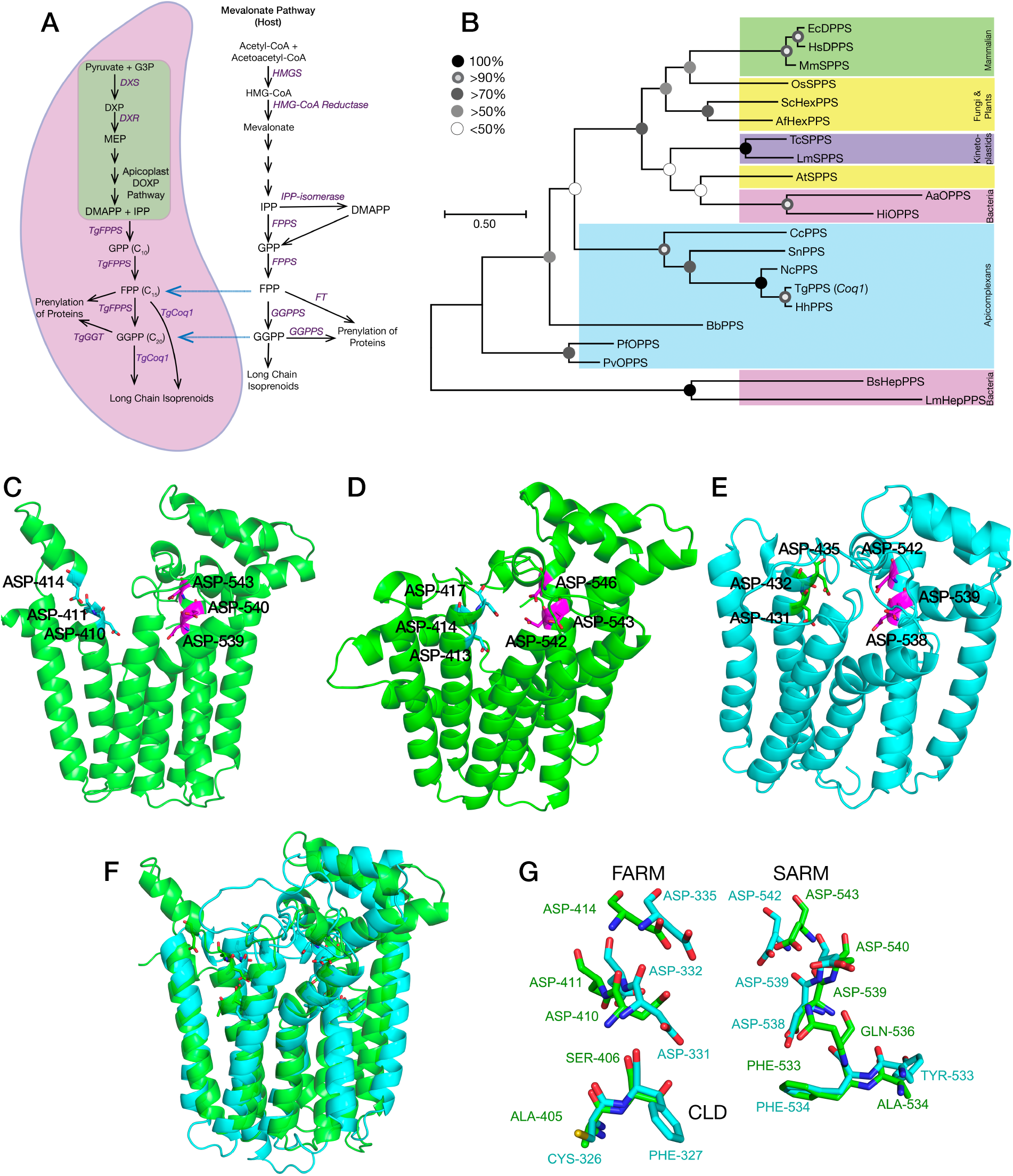
Phylogenetic placing and predicted structure of the *Toxoplasma* polyprenyl synthase (TgCoq1) and its predicted role in the isoprenoid pathway. **A**, the isoprenoid (DOXP/MEP) pathway of *T. gondii* compared to the host Mevalonate (MVA) pathway. G3P, glyceraldehyde 3-phosphate; DXP, deoxyxylulose 5-phosphate; DXS, DXP synthase; DXR, deoxyxylulose 5-phosphate reductoisomerase; MEP, methyl-D-erythritol phosphate; DMAPP, dimethyl ally diphosphate; IPP, isopentenyl diphosphate; GPP, geranyl diphosphate; FPP, farnesyl diphosphate; FPPS, farnesyl diphosphate synthase; GGPP, geranylgeranyl diphosphate; GGT, geranylgeranyl transferase; HMGS, 3-hydroxy-3-methylglutaryl-CoA synthase; FT, farnesyl transferase; GGPPS, geranylgeranyl diphosphate synthase. The *blue arrows* represent the intermediate metabolites that can be salvaged from the host. **B**, Phylogenetic analysis of enzymes of the isoprenoid pathway from various organisms. The sequences used are listed in supplemental Table S1. The enzymes are HexPPS, hexaprenyl pyrophosphate synthase; HepPPS, heptaprenyl pyrophosphate synthase; OPPS, octaprenyl pyrophosphate synthase; SPPS, solanesyl pyrophosphate synthase; DPPS, decaprenyl pyrophosphate synthase and PPS, polyprenyl synthase. The analysis shows substitutions per site for *Arabidopsis thaliana* SPPS (NP_17792.2), *Oryza sativa* SPPS (Q653T6.1), *Saccharomyces cerevisiae* HexPPS (P18900.1), *Trypanosoma cruzi* SPPS (EAN82722.1), *Leishmania major* SPPS (XP_001682016.1), *Plasmodium falciparum* OPPS (XP_001349541.1), *Plasmodium vivax* OPPS (VUZ93874.1), *Toxoplasma gondii* polyprenyl synthase (TgCoq1) trimmed (XP_018636584.1), *Neospora caninum* PPS (XP_003883948.1), *Hammondia hammondi* PPS trimmed (HHA_269430), *Cyclospora cayetanensis* PPS (LOC34620908), *Sarcocystis neurona* PPS trimmed (SN3_03700020), *Babesia bovis* PPS (EDO07087.1), *Bacillus subtilis* HPPPS (ARW31988.1), *Listeria monocytogenes* HepPPS (QGK56962.1), *Aquifex aeolicus* OPPS (NP_213606.1), *Haemophilus influenzae* HepPPS (CBY85984.1), *Aspergillus fumigatus* DPPS (EDP53123.1), *Mus musculus* SPPS (BAE48219.1), *Equus caballus* DPPS (XP023487828.1), and *Homo sapiens* DPPS (BAE48216.1). **C,** Phyre2 prediction based on the full TgCoq1sequence: only the catalytic domain is predicted, and it is as found in other *trans*- head-to-head prenylsynthases. The catalytic Asps in the two DDxxD motifs are indicated. **D**, Phyre2 prediction based on the full HhFPPS sequence. **E**, Phyre2 prediction based on the full TgFPPS sequence. **F**, Superposition of TgCoq1 (green) and TgFPPS (cyan) predicted structures. **G**, Zoomed-in version of F.

In this work we characterize the enzyme (*TgCoq1*) responsible for the synthesis of the isoprenoid chain that supplies the tail of the mitochondrial ubiquinone, as well as determine its biological function in the parasites. In addition, we show that specific inhibition of TgCoq1 impacts parasite growth, as well as its virulence in mice. Our results thus reveal that TgCoq1 is a promising chemotherapeutic target for the development of new medicines for toxoplasmosis.

## RESULTS

### Identification of *T. gondii* Coq1

The *T. gondii* gene *TGGT1_269430* annotated in ToxoDB as a polyprenyl synthase predicts the expression of a protein of 676 amino acids with a calculated molecular mass of 72.5 kDa and an isoelectric point of 8.54. The deduced *T. gondii* polypeptide sequence was aligned with other representative polyprenyl diphosphate synthases (Fig. S1) to highlight the specific domains, in particular the first aspartic rich motif (FARM) and the second aspartic rich motif (SARM) involved in substrate recognition [14]; the chain-length-determining (CLD) region, as well as a long N-terminal sequence thought to be involved in mitochondrial targeting. All the conserved motifs involved in catalysis or binding (regions I–VII) identified in other polyprenyl synthases (PPS) [16] are present in the *T. gondii* enzyme (Fig. S1). The presence of small residues (Ser and Ala) at positions 4 and 5 prior to the first aspartate-rich motif (FARM) predicts that the active site of the enzyme will accommodate long-chain isoprenoid products [17]. These residues are Phe in human FPPS, which makes only the C15 compound FPP. That is, bulky amino acids would not permit the nascent isoprenoid chains to extend further inside the hydrophobic cavity of the enzyme. A BLAST search of the protein data base showed that the amino acid sequence from *T. gondii* shared 81.31% identity with *Neospora caninum* PPS, 92.21% identity with *Hammondia hammondi* PPS, and 29.98% identity with the *Plasmodium falciparum* octaprenyl diphosphate synthase (OPPS). For the human homologue (accession number NP_055132), the identity was 31.15% [18]. We named this gene *TgCoq1*, following the established nomenclature of ubiquinone biosynthesis enzymes [6].

A phylogenetic comparison of the TgCoq1 sequence with representative animal, plant, yeast and other apicomplexan long prenyl diphosphate synthases (Fig. 1B and Table S1) showed that TgCoq1 groups with long-chain isoprenoid-biosynthesis enzymes from other organisms and is most closely related to other apicomplexan enzymes known or predicted to be involved in the synthesis of long-chain prenyldiphosphates. Notably, the *T. gondii* Coq1 protein is predicted to have a very long N-terminal domain, which was not reported in earlier annotations of ToxoDB. We used multiple programs to analyze the mitochondrial targeting sequence prediction and cleavage site. TargetP 2.0 program predicts 0.94 and MitoProt II predicts 0.7689 probability of targeting to mitochondria. The MitoProt II program also predicts the presence of a cleavage site 54 aa from the N terminal. A similar N-terminal extension is found in the orthologous gene of *Hammondia hammondi*, an avirulent relative of *T. gondii,* but has not yet been reported in other apicomplexan sequences.

To delve more into the structure of TgCoq1 we next used the Phyre2 program [19] for structure prediction. Using a diverse range of templates, several similar structures were predicted with 100% confidence (with an average of ∼30% identity). However, in all cases, only approximately one-half of the protein structure—the catalytic C-terminal domain— was successfully predicted, as shown in Figure 1C. The predicted structure of the catalytic domain of the *H. hammondi* protein is quite similar and it is shown in Figure 1D. The alphafold server [20] also was unable to predict the N terminal.

Of note, a long N-terminal domain is also found in the previously characterized *T. gondii* FPPS (TgFPPS) [14], a protein with 646 residues, slightly smaller than TgCoq1 (676 residues). The Phyre2 structure prediction for TgFPPS is shown in Figure 1E and again, only the catalytic domain can be successfully predicted. Figure 1F shows a superposition of the predicted structures for TgCoq1 (green) and the TgFPPS (cyan) showing high similarity. Figure 1G shows a zoomed-in view of Figure 1F (same color scheme) with the Ser/Ala/Phe CLD region indicated. From these results and >100 previously reported FPPS structures, we concluded that the presence of the bulky residues FF at positions 4 and 5 upstream to the SARM are found exclusively in the enzymes that synthesize the C15 isoprenoid product (e.g. human FPPS), while the aminoacids CF are present in the bifunctional TgFPPS, which synthesizes C15 as well as C20. For the polyprenyl synthase, TgCoq1, the aminoacids at the same position are AS, enabling extensive chain elongation. However, the H-bond network seen in the neutron crystallographic structure of human FPPS and proposed to be involved in H^+^ transport in all *trans*-head-to-head prenylsynthases [21, 22] is also present in TgCoq1. Specifically for the HsFPPS/TgCoq1 comparison: Thr-201/Thr-499; Gln-240/Gln-536; Asp-243/Asp-539; Asp-244/Asp-540; Tyr-193/Tyr-491 are predicted to have similar interactions, as it is found in longer-chain species, such as hexaprenyl and heptaprenyl diphosphate synthases [23, 24].

We next considered possible ‘physical’ properties of the N-terminal regions in TgCoq1, as well as TgFPPS. We used the HeliQuest program [25] to investigate the hydrophobicity <**H**>, hydrophobic moment <**μH**) and charge in the N-terminal regions of both proteins (Figs. S2A, S2B and Table S2). The N-terminus of TgCoq1 appears to have a hydrophobic region (closest to the actual N-terminus), followed by a very polar cationic domain, and then an anionic domain. In sharp contrast, no such pattern was observed in the N-terminus of the TgFPPS (Fig. S2B), suggesting the possibility that the N-termini might have different targets. We show in Table S2 helical wheel representations of the 3 regions using an 18 residue α-helix scan in which the likely neutral/hydrophobic, cationic and then anionic regions, with net charges of 0, +5 and −5, are shown. These regions are not apparent in the sequence of the TgFPPS, which instead is highly enriched in serine residues [14]. There is only a ∼10 residues sheet secondary structure predicted, all other residues being either in helical or disordered regions.

In summary, the structures of the catalytic domains of TgCoq1 and TgFPPS are predicted to be very similar. However, both proteins have extended N-terminal domains, and with the exception of the polyprenyl synthase sequence of *Hammondia hammondi*, they are not found in any other polyprenyl synthase sequence. As discussed below, these domains are important for mitochondrial targeting and perhaps, for other functions that remain to be determined.

### TgCoq1 synthesizes heptaprenyl diphosphate and *T. gondii* produces ubiquinone 7 (UQ_7_)

To investigate the biochemical function of TgCoq1 we cloned the *TGGT1_269430* gene in an *E. coli* expression plasmid which also includes an N-terminal polyhistidine tag to facilitate the purification of the recombinant protein by affinity chromatography. The activity of the purified protein was determined using allylic substrates and co-substrates, by measuring the amount of IPPs incorporated into the different long chain polyprenols (FPP and GGPP) (Fig. 2A). The reaction product was dephosphorylated and analyzed by reverse-phase thin layer chromatography (TLC), which showed labeling of a C35 product (Fig. 2B), indicating that TgCoq1 synthesizes heptaprenyl diphosphate. A parallel experiment with the already characterized recombinant *Trypanosoma cruzi* solanesyl diphosphate synthase [26], TcSPPS, known to synthesize a 45-carbon product, is shown in the same TLC (Fig. 2B). Using a standard enzymatic method [26], the activity of TgCoq1 was shown to be more efficient with FPP than with GGPP (Fig. 2C) or GPP as substrate (Table 1). The kinetic parameters of the reaction using different substrates is shown in Table 1.

**Figure 2.**
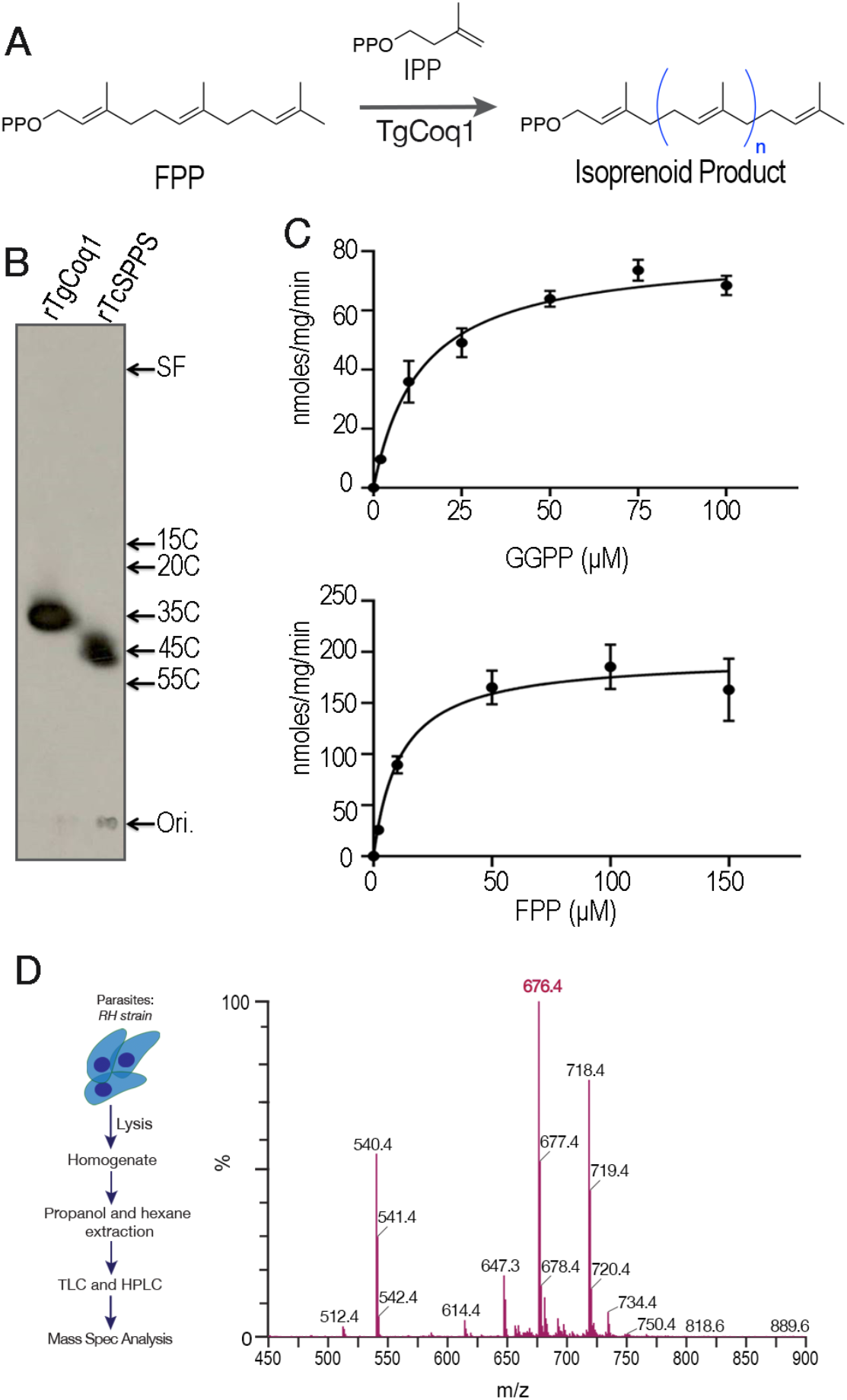
TgCoq1 catalyzes the synthesis of long chain isoprenoids from FPP. **A**, Scheme of the reaction catalyzed by Coq1. **B**, Analysis of the products formed by rTgCoq1 by thin-layer chromatography. The size of the product made by rTgCoq1 has 35 carbons (heptaprenyl). The reaction catalyzed by the *T. cruzi* solanesyl diphosphate synthase was run in parallel (rTcSPPS). Commercial standards of various chain lengths were run in the same system. SF is the solvent front and Ori. is the loading level of the samples. **C**, Specific activity assays of recombinant TgCoq1. Graphical data supporting the calculations of *K_m_* and *V_max_* for TgCoq1 are presented in Table 1. **D**, Mass spec analysis of *T. gondii* extracts obtained by 1-propanol followed by hexane extraction (see methods). Extracts were first purified by TLC, followed by HPLC prior to mass spec.

**Table 1:** Enzymatic Characterization of recombinant TgCoq1. Allylic substrate specificity of TgCoq1. The concentration of the allylic substrates (GPP, FPP, GGPP and IPP) was varied, and the counter substrate concentration was kept at a saturating level. The nonlinear regression analysis (Sigma plot 10.0) was used to calculate the Kinetic parameters. Values shown are means ± SD of two independent experiments performed in duplicate. The values of *k_cat_/K_m_* show that the enzyme is more efficient with FPP as the allylic substrate.

| Substrate | Counter Substrate | $K_m$<br>(μMolar) | $V_{max}$<br>(nmoles/min/mg) | $K_{cat}$<br>($\text{min}^{-1}$) | $K_{cat}/K_m$<br>($\text{min}^{-1} \times \text{Mole}^{-1}$) |
| --- | --- | --- | --- | --- | --- |
| GPP | IPP (100μM) | $0.88 \pm 0.60$ | $6.16 \pm 0.45$ | $0.012 \pm 0.001$ | 13.63 |
| FPP | IPP (100μM) | $12.37 \pm 1.83$ | $226.13 \pm 7.56$ | $0.452 \pm 0.015$ | 36.54 |
| GGPP | IPP (100μM) | $14.05 \pm 1.43$ | $82.98 \pm 2.51$ | $0.165 \pm 0.005$ | 11.74 |
| IPP | GPP (150μM) | $145.48 \pm 1.18$ | $28.30 \pm 0.16$ | $0.057 \pm 0.001$ | 0.39 |
| IPP | FPP (100μM) | $139.11 \pm 17.68$ | $560.54 \pm 46.09$ | $1.12 \pm 0.092$ | 8.05 |
| IPP | GGPP (50μM) | $141.89 \pm 35.75$ | $372.92 \pm 65.53$ | $0.745 \pm 0.131$ | 5.25 |

Since the length of the isoprenoid unit synthesized by Coq1 defines the type of UQ found in cells, we next extracted freshly purified tachyzoites and analyzed the extracts by TLC followed by HPLC and mass spectrometry (Fig. 2D). Standards for UQ_6_, UQ_7_, UQ_8_, UQ_9_, and UQ_10_ were run and the retention time for the ubiquinone isolated from *T. gondii* coincides with that of UQ_7_. The MS spectra shown in Fig. 2D highlights the specific m/z value of UQ_7_ in the form of ammonium adduction. The amount of UQ_7_ was quantified and found to be 4.2 µg/10^10^ tachyzoites (Fig. S3A).

We further validated the heptaprenyl diphosphate synthesis activity of TgCoq1 by expressing the gene in *E. coli* and extracting UQ (Fig. S3B and C). As shown in Fig. S3C, the synthesis of UQ_7_ was enhanced after bacteria was induced to express TgCoq1, indicating TgCoq1 is functional in *E. coli* without requiring another subunit while the fission yeast and the human decaprenyl diphosphate synthase enzymes required two subunits for functionality [18, 27].

In summary, TgCoq1 synthesizes heptaprenyl diphosphate, a C35 species, and *T. gondii* makes UQ_7_, supporting the role of TgCoq1 as heptaprenyl diphosphate synthase.

### TgCoq1 is essential for *T. gondii* growth and localizes to the mitochondria

To investigate the subcellular localization of TgCoq1 we introduced a triple HA (human influenza hemagglutinin) epitope-tag at the 3’ end of the *TgCoq1* gene locus and isolated *TgCoq1-HA* clones resistant to chloramphenicol (Fig. 3A). We found that TgCoq1 co-localizes with the mitochondrial marker MitoTracker (Fig. 3B). We next modified the 5’ region of the *TgCoq1-HA* mutant inserting a tetracycline (ATc)-regulatable element [28] and isolated the *i*Δ*Coq1-HA* mutant in which the expression of the tagged TgCoq1 is controlled with anhydrotetracycline, ATc (Fig. 3A). The parental cell line was the *TatiΔku80* which combines regulated gene expression [28] with higher efficiency of homologous recombination [29]. We complemented the *i*Δ*Coq1-HA* mutant with the *TgCoq1* cDNA (*i*Δ*Coq1-coq1*) or with a cosmid (PSBME30) (*i*Δ*Coq1-cosmid*) containing the *TgCoq1* genomic locus. These genetic modifications generated the mutants *i*Δ*Coq1-HA, i*Δ*Coq1-coq1* and *i*Δ*Coq1-cosmid* which were validated by Southern blot analysis (Fig. S4A). Immunofluorescence analysis (IFA) showed that TgCoq1-3HA was not expressed after 3 days of culture with ATc (Fig. 3B, *bottom row*). Western blots showed that the expression of TgCoq1 was fully ablated at day 3 with ATc and significantly reduced at day 2 (Fig. 3C). We next assessed growth of the mutant using plaque assays in which the parasite engages in repetitive cycles of invasion, replication, and egress, lysing the host monolayer and forming plaques that can be visualized by staining cultures with crystal violet. Preincubation of the mutant with ATc for 3 days prevented the formation of plaques in the *i*Δ*Coq1* cell line and this growth defect was restored in the complemented *i*Δ*Coq1-coq1* mutant (Fig. 3D-E). To further characterize the growth defect of the *iΔCoq1* mutant we expressed a cytosolic fluorescent protein (tdTomato) in parental and mutant (*iΔCoq1*) lines and used its fluorescence as a proxy for growth of clonal lines [30] (Fig. 3F). The growth defect of the *iΔCoq1* mutant was partially rescued by expression of Coq1 in the *iΔCoq1-coq1* mutant (Fig. 3F). To further characterize the growth defect, we tallied the number of parasites per parasitophorous vacuole (PV) 24 h post-infection and found that most PVs of the *i*Δ*TgCoq1*(+ATc) contained 2 parasites, some with 4 parasites and none with 8 or 16. Under identical conditions, the parental and the complemented mutants showed a similar distribution pattern with PVs containing 8 and 16 parasites (Fig. 3G). We next tested if removing ATc would result in recovery of growth of the mutant. With this aim, we grew the *iΔCoq1* mutant with ATc for 7 days and then removed the ATc and let the parasites grow for an additional 7 days and then measured the size of plaques. We found that the growth defect from the loss of coq1 could be irreversible as the plaques were significantly smaller as the control without ATc lysed the monolayer in 14 days and were unquantifiable. This partial reversibility would be expected as it is a knockdown cell line and not a clean knockout mutant (Fig S4D). These results support an essential role for TgCoq1 in intracellular *T. gondii* replication.

**Figure 3.**
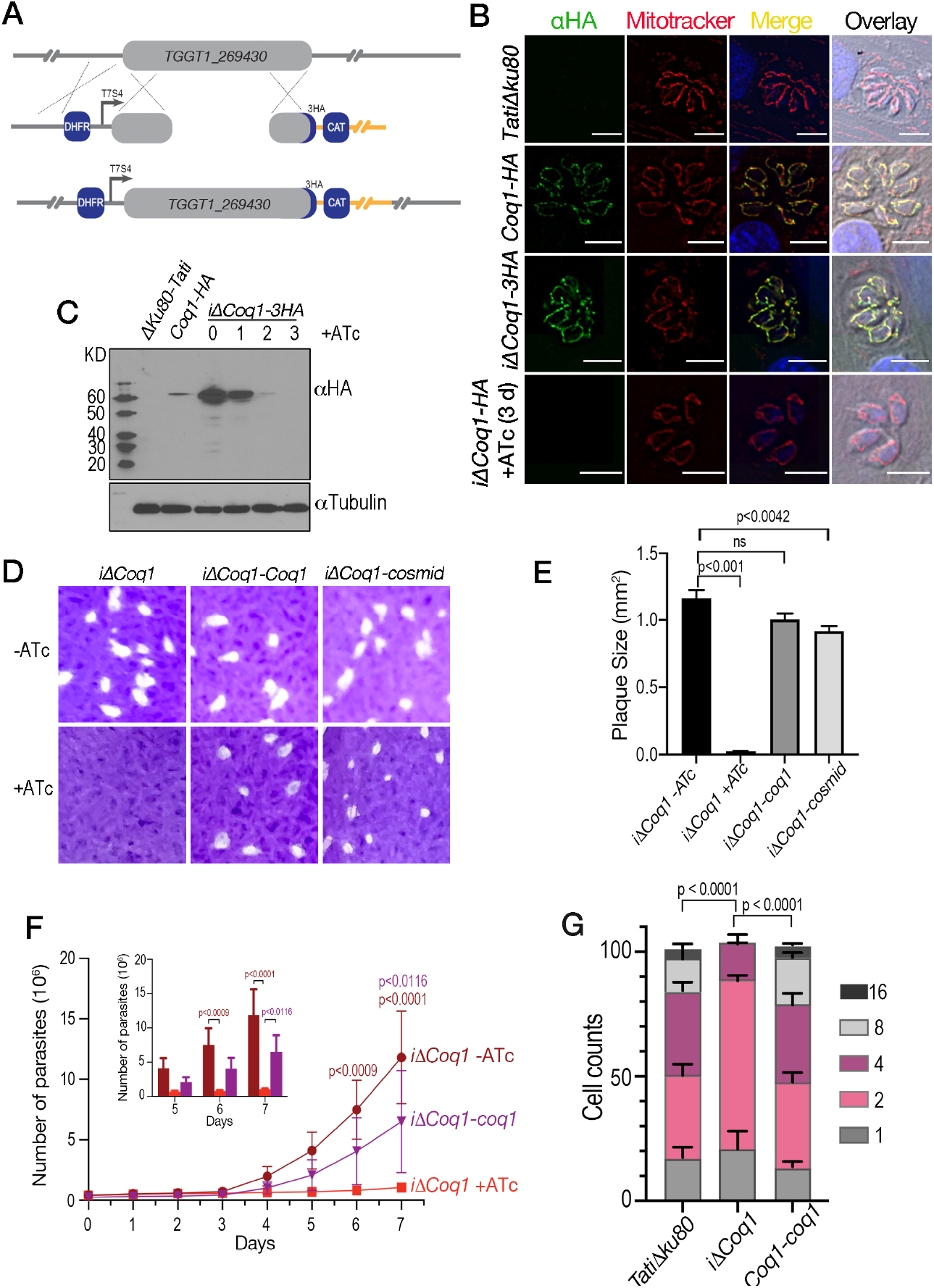
TgCoq1 is essential for *T. gondii* growth and localizes to the mitochondria. **A**, Strategy used to tag the *TgCoq1* gene locus with 3xHA and promoter replacement strategy for generation of the *iΔCoq1-3xHA* mutant. The endogenous TgCoq1 promoter is replaced with a Tet7Sag4 (T7S4) tetracycline inducible promoter. Expression of the dihydrofolate reductase (DHFR) confers pyrimethamine resistance and was used as a selectable marker. **B**, Immunofluorescence showing TgCoq1 localization to the mitochondria. Mitotracker was used to stain the mitochondria, and anti-HA antibody to localize TgCoq1. The TgCoq1 signal in the *iΔCoq1-3xHA* mutant at 0 and after 3 days of growth with 0.5 µg/ml ATc. **D,** Plaque assay showing that the *iΔCoq1* mutant was unable to form plaques when grown with ATc. Parasites were pre-incubated with ATc for 3 days prior to the experiment. Growth was restored by complementation with *TgCoq1* cDNA or with a cosmid containing the full *TgCoq1* genomic locus. **E**, Plaque size quantification from three biological experiments. **F**, Growth assay of RFP expressing cells highlighting the growth defect of the *iΔCoq1* mutant and its rescue by complementation with TgCoq1 cDNA. **G**, Parasite replication assay showing the parasites/vacuole for each of the strains. Analysis was performed from three independent biological experiments, each one in triplicate. Statistical analysis for E, F and G was done using Student’s t-test.

### A different isoprenoid tail can replace the one synthesized by TgCoq1

The *Trypanosoma cruzi* solanesyl diphosphate synthase (TcSPPS) was previously characterized and shown to synthesize a product with 9 isoprene units (C45) [26]. We wondered if the length of the isoprenoid chain was important for the biological function of UQ in *T. gondii* so we next complemented the *i*Δ*Coq1* mutant with the *TcSPPS* gene (Fig. 4A). Since TgCoq1 localizes to the mitochondrion, we created a construct with the sequence of the N-terminal signal peptide from TgCoq1 and inserted it upstream to the *TcSPPS* gene (Fig. 4A). Southern blot analysis confirmed the presence of the *TcSPPS* gene in the *i*Δ*Coq1*-*tcspps* mutant (Fig. S4B). Western blot analysis with αHA antibodies show a band at 58 kDa which disappears upon ATc treatment for 3 days. This band also decreased in cells complemented with *TcSPPS* upon treatment with ATc, while a band that could correspond to the TcSPPS (56 kDa) did not (Fig. S4C). Further validation for the downregulation of TgCoq1 is shown in the western blots using anti-TgCoq1 antibody generated against the whole recombinant protein (Fig. 4B). The 58 kDa band is absent in the *iΔCoq1* mutant after culturing with ATc, while it is present in the complemented mutant (grown with and without ATc). The anti-TgCoq1 antibody did not react against the TcSPPS and it did not show a reaction in the *iΔCoq1-tcspps* (+ATc). The IFAs in Fig. 4C showed the localization of TcSPPS to the mitochondria of the *iΔCoq1-tcspps* mutant, as detected by co-localization with antibodies against TOM40 [31].

**Figure 4.**
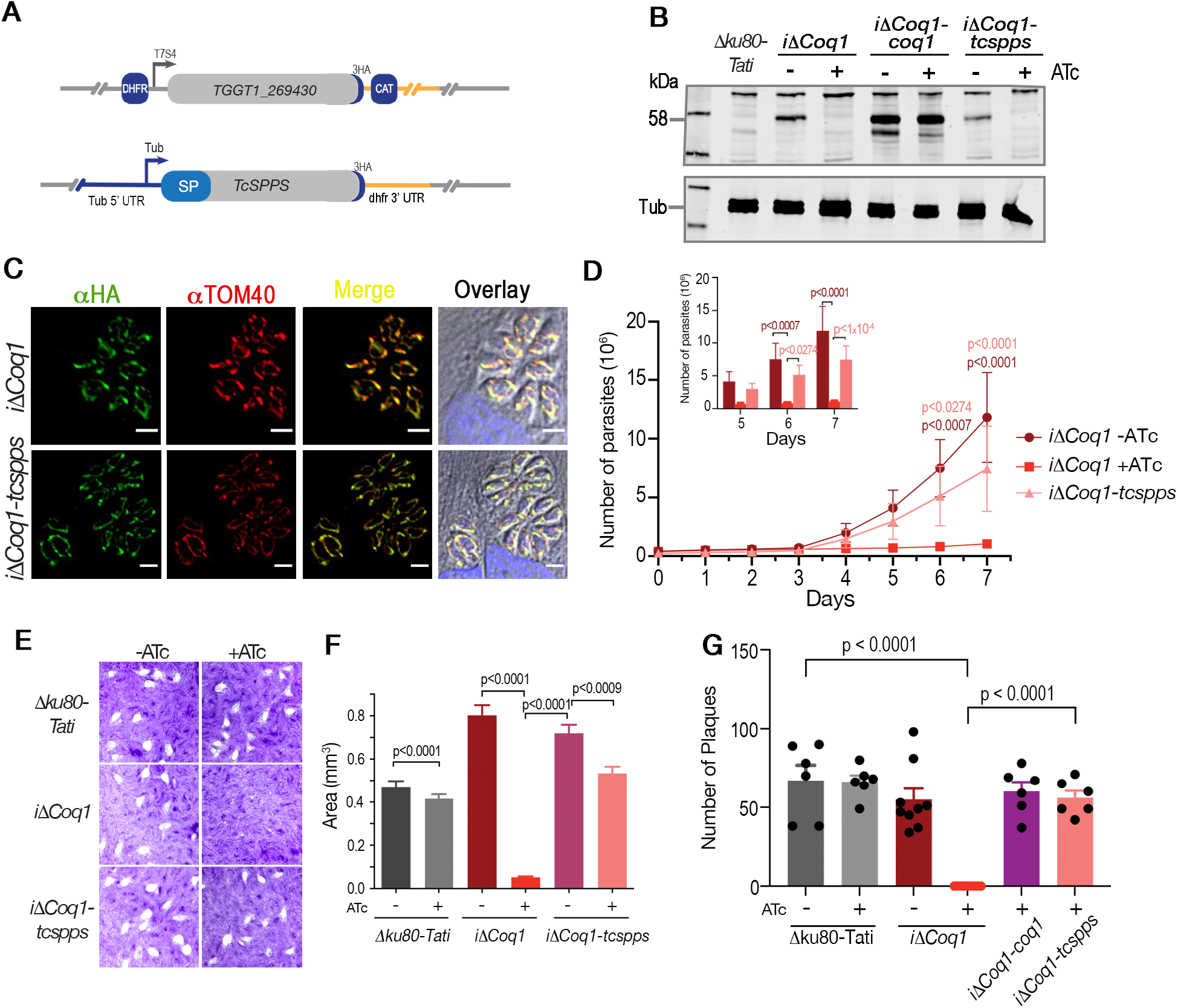
A different isoprenoid tail can replace the one synthesized by TgCoq1. **A**, Model depicting the complementation of the promoter insertion cells line (*i11Coq1*) with a *T. cruzi* SPPS in the pCTH3 plasmid backbone. **B**, Western blots with the polyclonal antibody against TgCoq1 (αTgCoq1) showing that the signal of TgCoq1 disappears in cells cultured with ATc. Tubulin was used for loading control. **C**, IFAs with αHA showing the localization of TcSPPS to the mitochondria of intracellular parasites. *i11Coq1-spps* cells were continuously cultured with ATc and as shown in Fig. 3C there is no expression of the parental Coq1. **D**, Growth assay of RFP expressing cell lines showing that the growth defect of the *i11Coq1* mutant is restored by complementation with the *T. cruzi* SPPS. **E**, Plaque assay showing that the *i11Coq1* mutant was unable to form plaques when cultured with ATc, and growth was restored by expression of TcSPPS. **F,** Quantification of the area of fifteen plaques from each of 3 assays like the one shown in E. **G,** Plaquing efficiency assay in which plaque numbers were quantified from 3 biological replicates. The analyses presented in D, F and G were from three independent biological experiments using Two-way anova analysis.

The severe growth phenotype of the *i*Δ*Coq1(+ATc)* (∼5 % of control at 7d +ATc) was reverted by complementation of the mutant with TcSPPS (Fig. 4D), supporting equivalent function of the *T. cruzi* enzyme to TgCoq1 (Fig. 4E-F). A plaquing efficiency assay in which contact between parasites and host cells was limited to 30 min and the number of plaques enumerated (Fig. 4G) showed that TcSPPS expression rescued the phenotype of the *i*Δ*Coq1(+ATc)* mutant. Both plaque size and number were restored by complementation with the TcSPPS enzyme (Fig. 4G).

Collectively, these data showed that TgCoq1 is essential for *T. gondii* growth, and the length of the isoprenoid unit is not critical for its biological function.

### TgCoq1 is essential for mitochondrial function

The mitochondrion of *T. gondii* is critical for replication and UQ is essential for the function of the mitochondrial ETC. Considering that UQ is attached to the mitochondrial membrane through the isoprenoid unit synthesized by Coq1, we next investigated whether the *i*Δ*TgCoq1* (+ATc) mutant was unable to grow due to mitochondrial malfunction. We tested two important mitochondrial features: Oxygen consumption rate (OCR) and mitochondrial membrane potential (Fig. 5). We measured the rate of oxygen consumption under basal (state 2); ADP stimulated (state 3, oxidative phosphorylation); oligomycin-inhibited (state 4, minimum oxidative phosphorylation), and FCCP-stimulated (state 3u, uncoupled) conditions using digitonin-permeabilized parasites suspended in a buffer containing succinate as substrate (Fig. 5A, see scheme in Fig. S5). Mitochondrial activity of the *i*Δ*TgCoq1* (+ATc) mutant showed significantly lower basal OCR, Ox. Phos (+ADP) and min. Ox. Phos. (+ oligomycin) (Fig. 5B-C). Interestingly, respiration of the TcSPPS-complemented mutant was significantly higher, which could be the result of either higher expression or activity of TcSPPS, or both.

**Figure 5.**
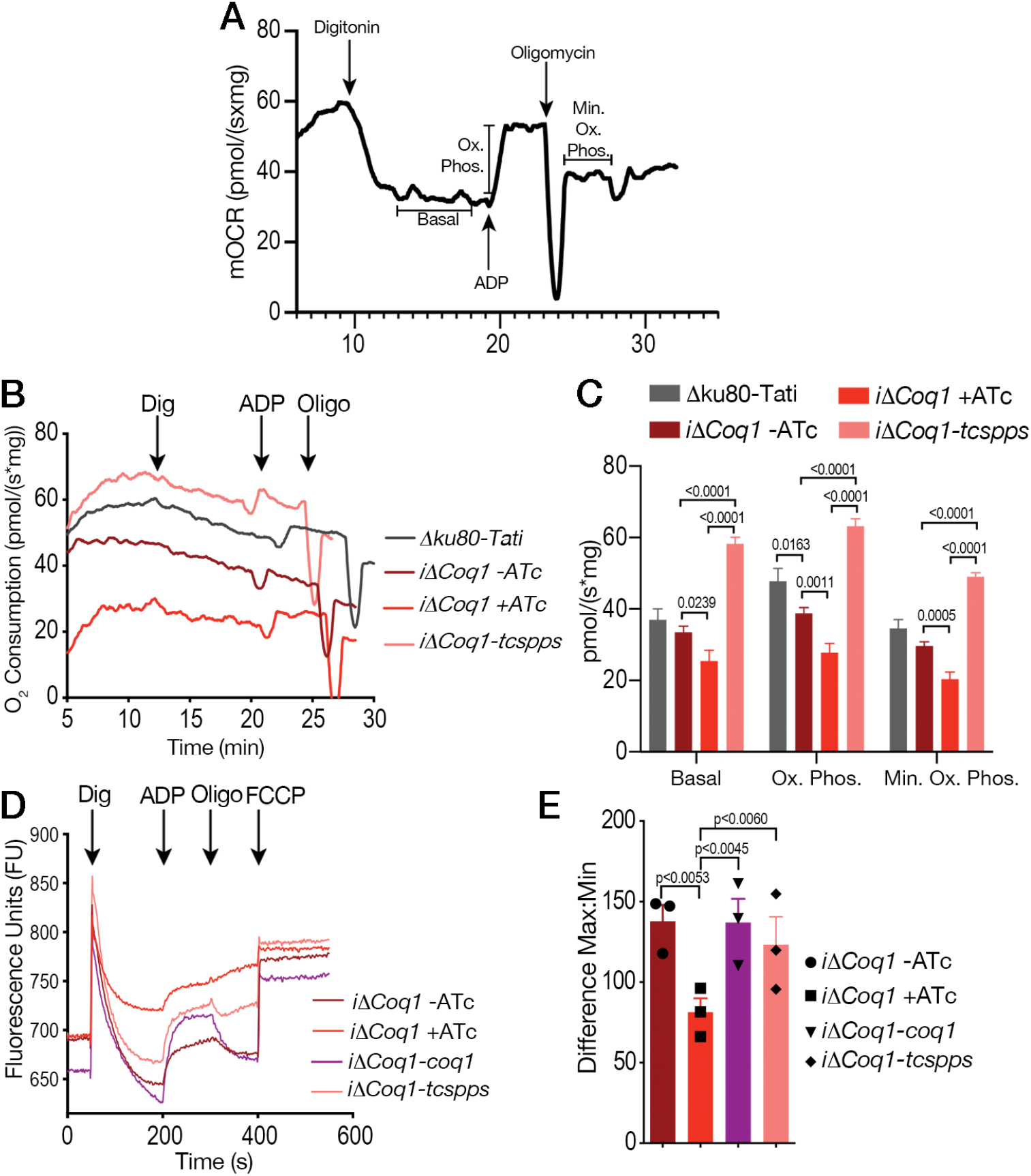
TgCoq1 is essential for mitochondrial function. **A,** Oxygen consumption rate (OCR) of RH*Δku80* parasites showing the computed parameters of each condition. **B,** OCR experiments measuring oxygen consumption of the *iΔCoq1* mutant showing a reduction in oxygen consumption after 3 days of incubation with 0.5 µg/ml ATc that is restored in the *T. cruzi* SPPS complemented cell lines also grown with 0.5 µg/ml ATc for 3 days. **C,** Quantification of peaks from the OCR experiments showing significant differences of the parental cell lines, *iΔCoq1*, and the *iΔCoq1-tcspps* complemented ones. **D,** Mitochondrial membrane potential (ΔΨm) of the *iΔCoq1* mutant without ATc and with 0.5 µg/ml ATc for 3 days. Defect in ΔΨm of the *iΔCoq1* mutant was detected after 3 days with ATc. Complementation with the TcSPPS gene restored the defect. Digitonin (Dig) was added at 50 sec, Adenosine diphosphate (ADP) was added at 200 sec, Oligomycin (Oligo) was added at 300 sec, and Carbonyl cyanide-4- (trifluoromethoxy) phenylhydrazone (FCCP) was added at 400 sec. **E,** Quantification of the mitochondrial membrane potential, the difference was calculated by subtracting the minimum after digitonin from the maximum peak after addition of FCCP. The dots represent each of 3 biological replicates which are the averages from 3 technical replicates. The analyses were done from three independent biological experiments using Student’s t-test.

We further characterized mitochondrial function by measuring the membrane potential (ΔΨ_m_) of digitonin-permeabilized tachyzoites (controls and mutants) in the presence of succinate as substrate, using safranine O [32]. Upon digitonin permeabilization, safranine O is inserted into energized mitochondrial membranes and causes fluorescence quenching. Once a steady state is reached, addition of ADP stimulates synthesis of ATP by the ATP synthase, which uses the H^+^ gradient, causing a partial depolarization of the membrane and release of safranine O (Fig. 5D). The fluorescence returned to its basal level upon addition of oligomycin, which inhibits ATP synthase activity and prevents further depolarization by ADP. FCCP addition uncoupled oxidative phosphorylation releasing safranine O (Fig. 5D). The *iΔCoq1* (+ATc) mutant was unable to load safranine up to the same level as the controls, indicating partial depolarization of the mitochondrial membrane potential (Fig. 5D and E). Complementation with TgCoq1 or TcSPPS restored mitochondrial membrane potential to the same level as the parental controls.

These results showed that TgCoq1 is essential for mitochondrial function in *T. gondii*, which was restored after complementation.

### The lipophilic bisphosphonate BPH-1218 inhibits TgCoq1 and protects mice against *T. gondii* infection

Previous work has shown that bisphosphonates can target several enzymes of the isoprenoid biosynthesis pathway, including the farnesyl diphosphate synthase (FPPS) which synthesizes FPP, the preferred substrate of TgCoq1 [14]. We screened for potential inhibitors that could specifically inhibit TgCoq1 and tested a series of ∼200 bisphosphonates. We measured the EC_50_s for the most promising candidates and in parallel we evaluated the inhibition of the recombinant TgCoq1 and recombinant TgFPPS, for comparison (Table 2). One interesting compound (BPH-1218) was more effective at inhibiting TgCoq1 than TgFPPS, with an IC_50_ of 36 nM for the inhibition of TgCoq1 and an EC_50_ of 0.92 µM against *T. gondii* growth *in vitro* (Table 2). We further determined specificity against TgCoq1, by comparing the growth inhibition of the compound against a parental strain (RH) versus a cell line over-expressing TgCoq1 (*TgCoq1-OE*). BPH-1218 (Fig. 6A), appears to specifically target TgCoq1, as supported by the 3-fold increase in EC_50_ against *TgCoq1-OE* compared to the parental strain (Fig. 6B, and Table 3). BPH-1218 was shown before to inhibit the TcSPPS with an IC_50_ of 60 nM and cause a 40% decrease in UQ_9_ biosynthesis in *T. cruzi* [33]. BPH-1218 showed very little toxicity when tested against human fibroblasts cells (Supplementary Table S3). We next tested if removing BPH-1218 would result in recovery of growth. We let parasites of the RH parental strain form plaques for 7 days in the presence of 1 μM or 2 μM BPH-1218. After 7 days we washed off the BPH-1218 and allowed the parasites to grow for 7 additional days undisturbed. The control wells were unquantifiable after 14 days as the host cell monolayer was completely lysed. However, the wells with BPH1218 showed plaques that did not increase in size and we saw that the plaque size of the 2 μM BPH-1218 were significantly smaller compared to the 1 μM (Fig 6D). This result indicates that the inhibition appears to be irreversible as we used concentrations close to the EC50 values of the drug and will only inhibit approximately 50% of growth.

**Figure 6.**
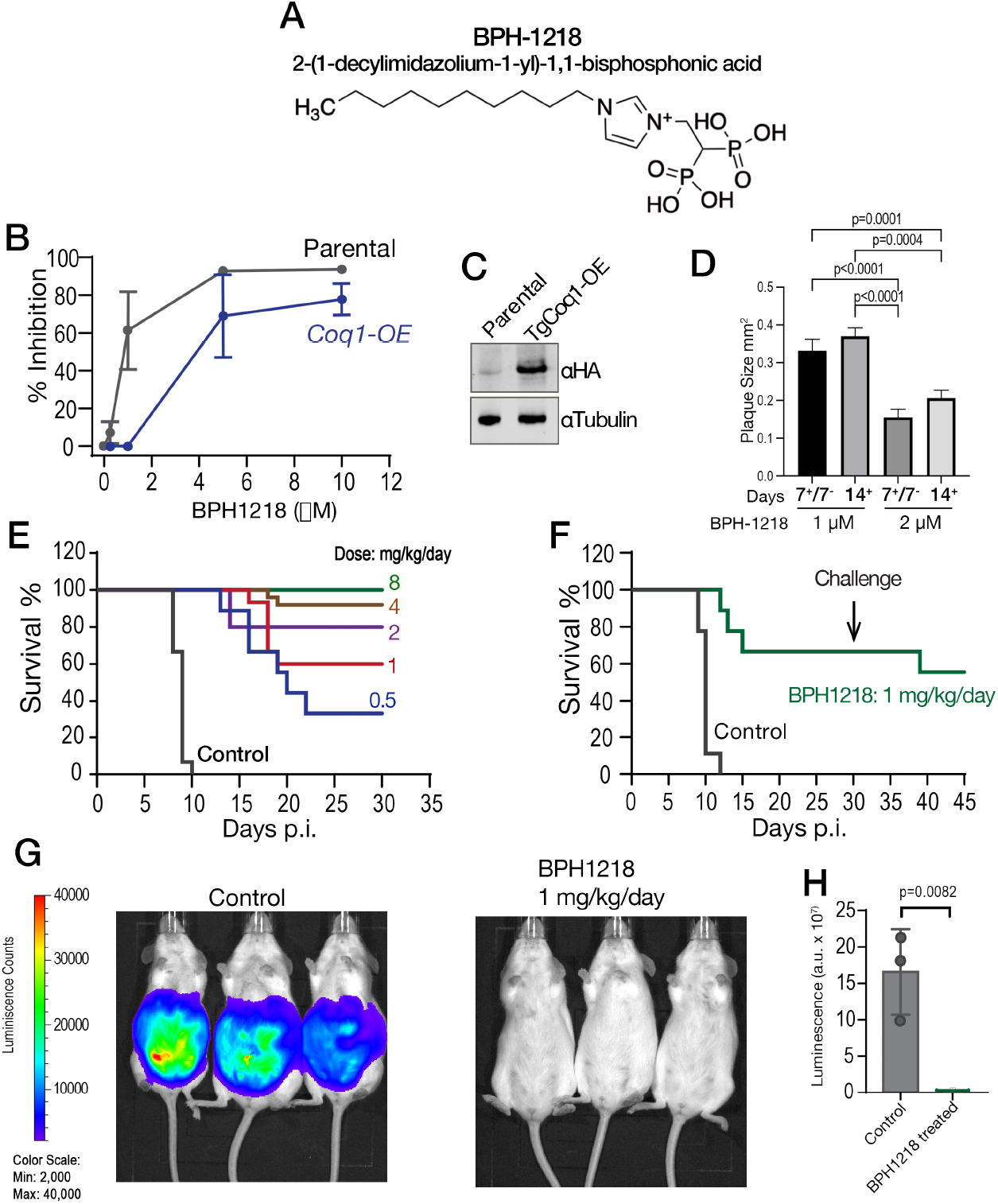
Targeting TgCoq1 with a lipophilic bisphosphonate. **A,** Structure of BPH-1218. **B,** % inhibition by the drug BPH-1218 of the parental and the TgCoq1 over-expressing mutant (Coq1-OE) showing that the Coq1-OE is more resistant to BPH-1218 (average from 3 independent experiments). **C,** Western blot analysis showing the increase in expression of the HA tagged TgCoq1 in the overexpressing cell line. **D,** Quantification of plaque sizes for growth of the wild-type cell line grown with BPH-1218 for 7 days and then without drug for 7 days or continuously grown in BPH-1218 for 14 days (BPH-1218 at indicated concentrations). We also tested no BPH-1218 and the parasites completely lysed the monolayer and not quantifiable (data summarized from 3 biological replicates, Two-way anova statistical analysis). **E,** BPH-1218 protects against a lethal *in vivo* infection in mice. The indicated doses (mg/kg/day) of BPH-1218 were inoculated for 10 days after mice were infected with a lethal dose of RH parasites. ED_50_ = 0.69 mg/kg/day (data summarized from three independent experiments). **F,** Challenge with 100 RH parasites of living mice on day 30 after treatment for 10 days with 1 mg/kg/day BPH-1218. **G,** Infection of mice with a luciferase-expressing strain of RH parasites shows BPH treatment of mice clears the infection. More details in Materials and Methods. **H,** Quantification of luciferase expression from two experiments, each with six mice (3 controls and 3 treated).

**Table 2.**
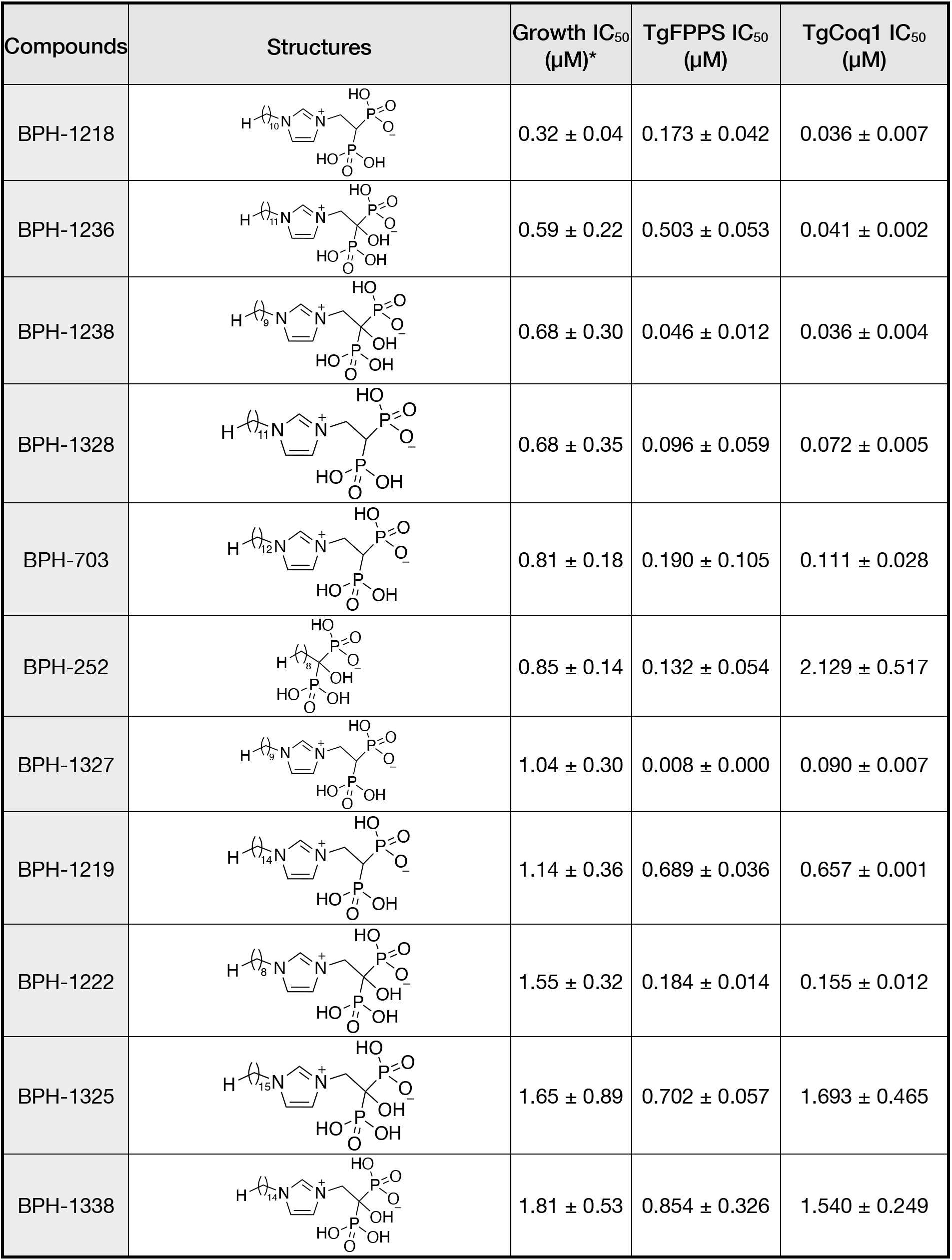
Inhibition of growth and the recombinant enzymes TgFPPS and TgCoq1 by selected bisphosphonates. * These compounds showed inhibition of at least 50% at 10 µM

| Compounds | Structures | Growth IC<sub>50</sub><br>(μM)* | TgFPPS IC<sub>50</sub><br>(μM) | TgCoq1 IC<sub>50</sub><br>(μM) |
| --- | --- | --- | --- | --- |
| BPH-1218 | | 0.32 ± 0.04 | 0.173 ± 0.042 | 0.036 ± 0.007 |
| BPH-1236 | | 0.59 ± 0.22 | 0.503 ± 0.053 | 0.041 ± 0.002 |
| BPH-1238 | | 0.68 ± 0.30 | 0.046 ± 0.012 | 0.036 ± 0.004 |
| BPH-1328 | | 0.68 ± 0.35 | 0.096 ± 0.059 | 0.072 ± 0.005 |
| BPH-703 | | 0.81 ± 0.18 | 0.190 ± 0.105 | 0.111 ± 0.028 |
| BPH-252 | | 0.85 ± 0.14 | 0.132 ± 0.054 | 2.129 ± 0.517 |
| BPH-1327 | | 1.04 ± 0.30 | 0.008 ± 0.000 | 0.090 ± 0.007 |
| BPH-1219 | | 1.14 ± 0.36 | 0.689 ± 0.036 | 0.657 ± 0.001 |
| BPH-1222 | | 1.55 ± 0.32 | 0.184 ± 0.014 | 0.155 ± 0.012 |
| BPH-1325 | | 1.65 ± 0.89 | 0.702 ± 0.057 | 1.693 ± 0.465 |
| BPH-1338 | | 1.81 ± 0.53 | 0.854 ± 0.326 | 1.540 ± 0.249 |

**Table 3:**
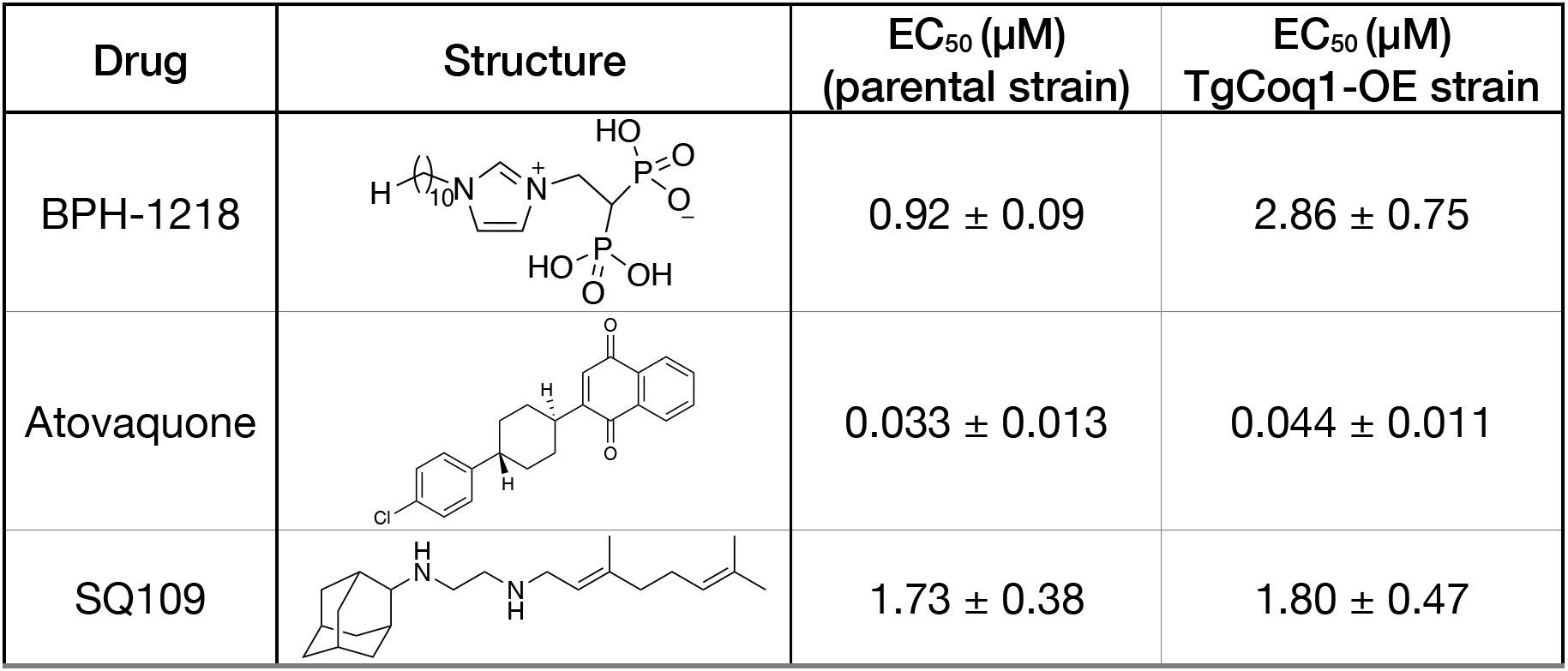
IC_50_ of BPH-1218 on TgCoq1-OE strain is significantly higher than parental strain while IC_50_ of drugs atovaquone and SQ-109 was not affected by TgCoq1 overexpression. Enzymatic activity shows that TgCoq1 is a more specific target than TgFPPS for BPH-1218.

| Drug | Structure | EC<sub>50</sub> (μM)<br>(parental strain) | EC<sub>50</sub> (μM)<br>TgCoq1-OE strain |
| --- | --- | --- | --- |
| BPH-1218 |  | 0.92 ± 0.09 | 2.86 ± 0.75 |
| Atovaquone |  | 0.033 ± 0.013 | 0.044 ± 0.011 |
| SQ109 |  | 1.73 ± 0.38 | 1.80 ± 0.47 |

We next tested BPH-1218 for in Swiss webster mice infected with 100 tachyzoites of the RH strain. We started treatment with BPH-1218 six hours post-infection and administered the drug daily for 10 days. We observed that BPH-1218 was protective against acute *T. gondii* infection *in vivo* with a ED_50_ of 0.69 mg/kg/day (Fig. 6E). The initial infection was successful as shown by the resistance to re-infection of the surviving mice (Fig. 6F). The protection by BPH-1218 was further shown in a model of dissemination of infection with Swiss Webster mice infected with a strain of *T. gondii* that expresses a luciferase gene (RH-cLuc-GFP, a gift from Jeroen Saiej). The treatment protocol was similar to the one described for the acute infection except that single daily dose of 1 mg/kg of weight were used for 8 days. At this point live mice were inoculated with δ-Luciferin for visualization of the infection in an IVIS illumination system [34]. Treatment with BPH-1218 showed a remarkable effect on both virulence and dissemination of the infection (Fig. 6G-H).

In summary, both *T. gondii* growth *in vitro* and mice virulence were blocked by pharmacological inhibition of TgCoq1.

### An UQ derivative rescues BPH-1218 growth inhibition and *iΔCoq1* (+ATc) growth defect

To further validate the role of TgCoq1 in the synthesis of ubiquinone we tested if supplementing cultures of the *iΔCoq1* (+ATc) mutant with various ubiquinones would rescue the growth defect (Fig. 7). Because UQ_7_ is not commercially available we tested UQ_6_, UQ_8_ and CoQ_10_ and observed a remarkable rescue of the growth of the *iΔCoq1*(+ATc) mutant by UQ_6_ (Fig. 7A). The other UQs did not have a significant effect and could be the result of poor solubility or permeability (Fig. 7B). We next wondered if UQ_6_ would rescue the growth inhibition by BPH-1218 (Fig. 7C-D). UQ_6_ was able to rescue both genetic and pharmacological inhibition of TgCoq1. We applied this strategy with a larger number of compounds and tested % growth of 18 known mitochondrial inhibitors (shown in Table S4) and their possible rescue by UQ_6_ (Fig. 7E, at 2x and 4x EC_50_s). We observed that for some compounds the % growth did not change by adding UQ_6_ to the culture but some of them showed a reduced inhibitory effect. We next seleced five compounds and determined their EC_50_ in the presence of UQ_6_, and we observed that for three of them (BPH-1218, BPH-1217 and BPH-1236) the EC_50_ was increased when the cultures were supplemented with UQ_6_ (Table 4). The growth inhibition by JAG21, a highly effective mitochondrial inhibitor [35], was not rescued by UQ_6_ but there was a modest rescue by the mitochondrial inhibitor atovaquone (Fig. 7E).

**Figure 7.**
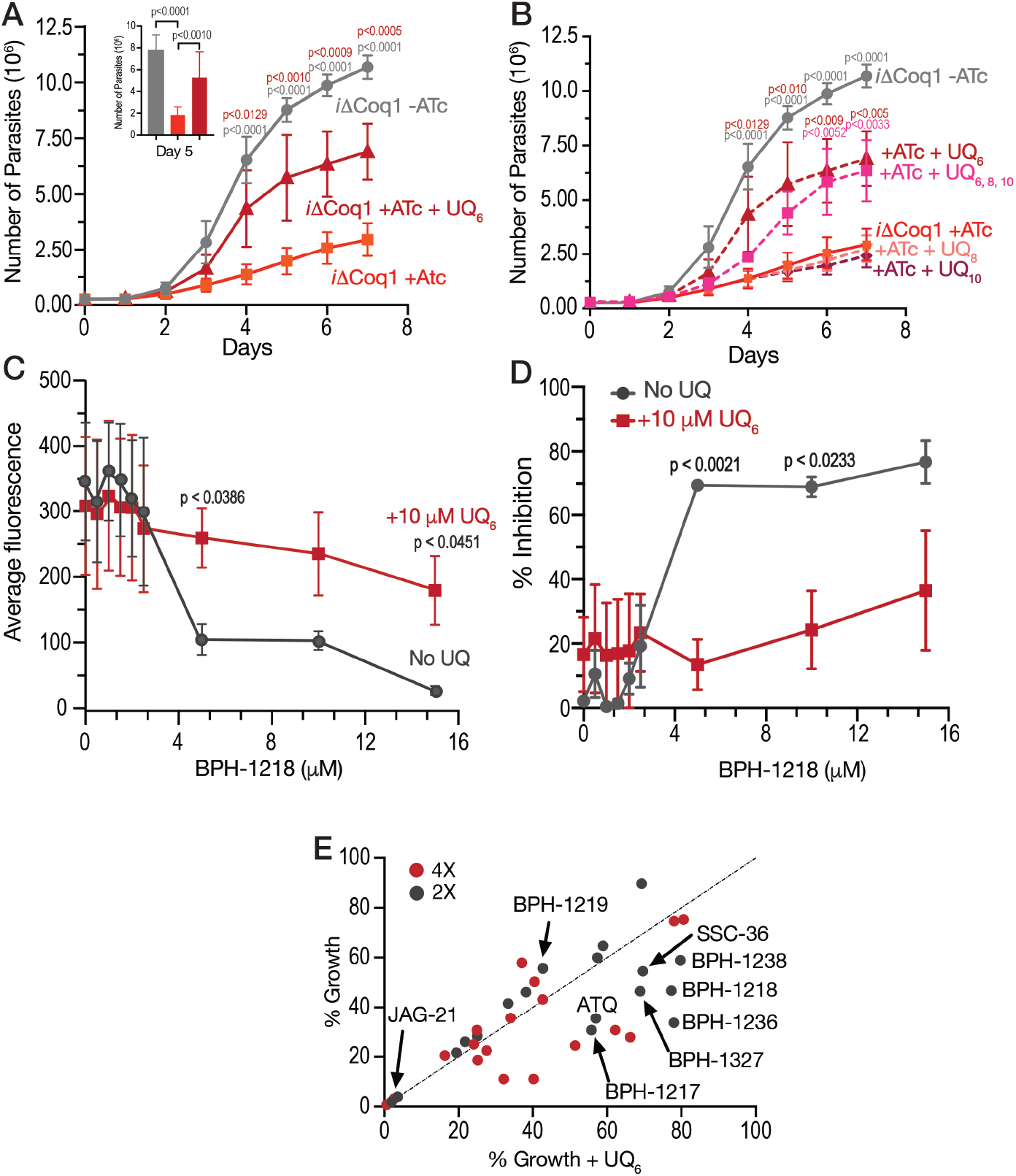
Ubiquinone 6 rescues the growth defect of the *iΔCoq1* mutant and the growth inhibition by BPH-1218. **A,** Growth of the *iΔCoq1* mutant with ATc is rescued by addition of 10 µM UQ_6_. **B,** Growth of the *iΔCoq1* with ATc supplemented with 5 µM Ubiquinone 8 (UQ8) and Ubiquinone 10 (UQ10) shows no rescue of the growth defect. Only UQ_6_ shows rescue at 5 µM. **C**, Inhibition by BPH-1218 is rescued by addition of 10 µM UQ_6_ to the culture. **D**, % inhibition by BPH-1218 with and without UQ_6._ (A-D: average from 3 biological replicates, Two-way anova statistical analysis). **E,** UQ_6_ rescue screen of 18 compounds at 2X and 4X their known EC_50_. Drugs that are rescued by UQ_6_ are shifted to the right from the dotted line. The dots are averages from 3 biological replicates. The compounds not labeled in the graph are BPH-1327, BPH-1238, Atovaquone, Pyrimethamine, BPH-754, Risedronate, SSC-36, CE-22, MNC-98, CE-29, MNCA181, CE-109, and CE-91. Structures shown in Table S5. (Average from 3 biological replicates).

**Table 4:**
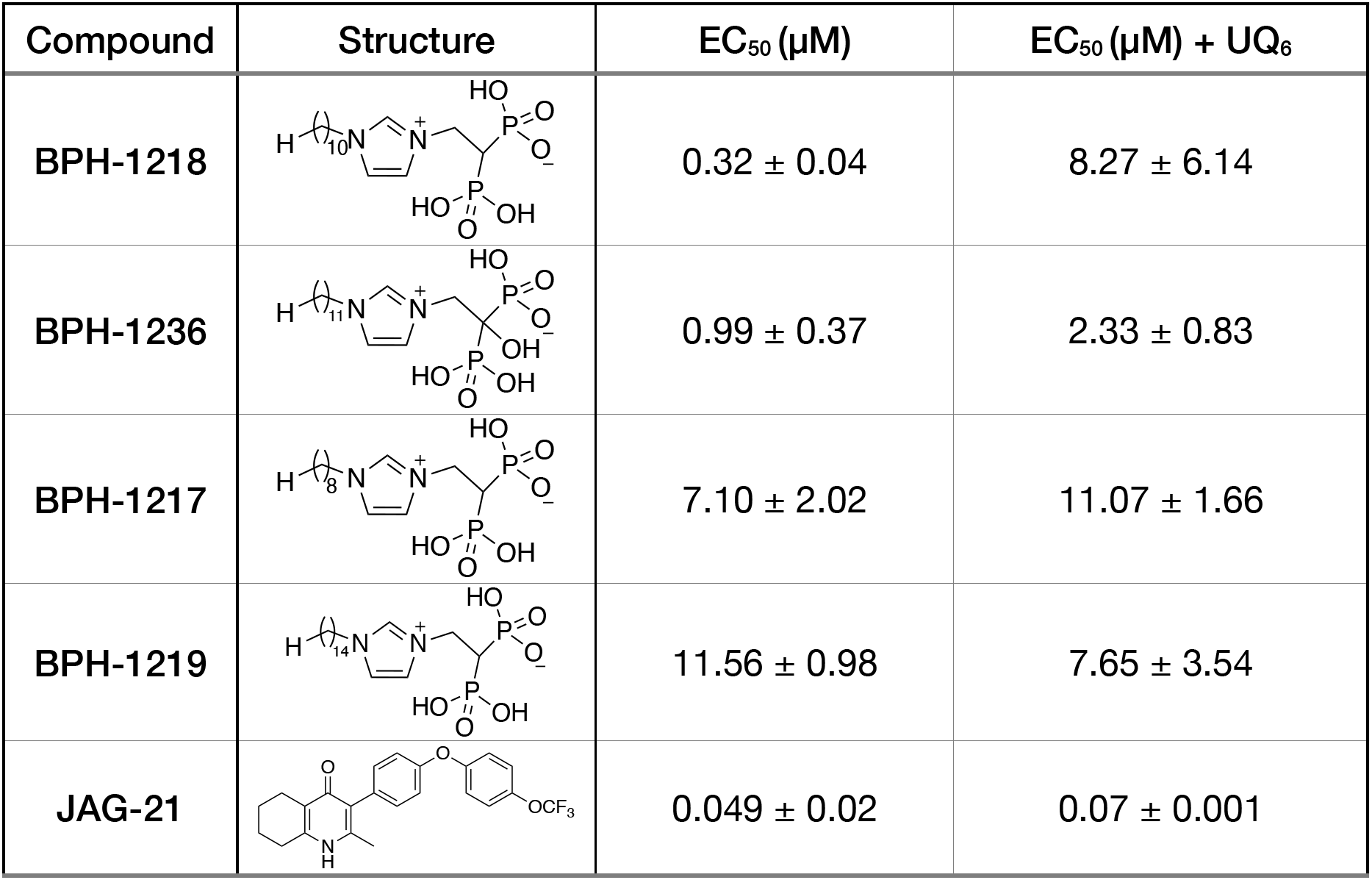
EC_50_ for *T. gondii* RH growth inhibition measured with and without the addition of ubiquinone Q_6_ to the culture. Values are means ± s.d. of n = 3.

| Compound | Structure | EC<sub>50</sub> (μM) | EC<sub>50</sub> (μM) + UQ<sub>6</sub> |
| --- | --- | --- | --- |
| BPH-1218 |  | 0.32 ± 0.04 | 8.27 ± 6.14 |
| BPH-1236 |  | 0.99 ± 0.37 | 2.33 ± 0.83 |
| BPH-1217 |  | 7.10 ± 2.02 | 11.07 ± 1.66 |
| BPH-1219 |  | 11.56 ± 0.98 | 7.65 ± 3.54 |
| JAG-21 |  | 0.049 ± 0.02 | 0.07 ± 0.001 |

In summary, TgCoq1 is an essential enzyme that forms part of the ubiquinone biosynthesis pathway and its pharmacological inhibition represents a new target that can be blocked with known bisphosphonates. Ubiquinone biosynthesis is an unexplored target in apicomplexan parasites and our work shows a strategy to discover new inhibitors via rescue with ubiquinones.

## DISCUSSION

In this work we report the characterization of a polyprenyl diphosphate synthase *TGGT1_269430* [36] (TgCoq1), the first enzyme of the ubiquinone synthesis pathway in *Toxoplasma gondii*. TgCoq1 localizes to the mitochondrion of *T. gondii* and is essential for parasite growth and infectivity. We showed that TgCoq1 synthesized heptaprenyl diphosphate (C35) and *T. gondii* produces UQ_7_. TgCoq1 was essential for normal mitochondrial function and both mitochondrial respiration and membrane potential were negatively impacted after genetically downregulating the expression of *TgCoq1*. We also show that over-expression of TgCoq1 protected parasite growth from the inhibitory effect of the lipophilic bisphosphonate BPH-1218. Supplementation with UQ_6_ rescued both the growth defect of the conditional mutant and the inhibition by the bisphosphonate. The bisphosphonate also protected mice infected with a lethal dose of *T. gondii* at an ED_50_ of 0.69 mg/kg/day.

Ubiquinones consist of a water-soluble (*p*-quinone/*p*-dihydroxyphenyl) headgroup that can accept or donate two electrons to the respiratory chain, and a lipophilic isoprenoid tail that targets the molecule to membranes. The isoprenoid tail is essential for the function of UQs in the electron transport chain of mitochondria [37]. The isoprenoid unit is derived from a common precursor, isopentenyl pyrophosphate (IPP), and its isomer, dimethylallyl pyrophosphate (DMAPP), which are synthesized in mammalian cells via the mevalonate pathway, but in *T. gondii* are synthesized via the apicoplast-localized methylerythritol phosphate (MEP) pathway (Fig. 1A), which is essential [7]. In *T. gondii*, IPP and DMAPP are condensed by the action of an unusual farnesyl diphosphate synthase (TgFPPS) into farnesyl diphosphate (FPP) and geranylgeranyl diphosphate (GGPP) [14]. Previous work from our group demonstrated that *T. gondii* imports FPP and GGPP, from the host cell, making them sensitive to host isoprenoid biosynthesis inhibitors, but at the same time less vulnerable to inhibitors of the parasite enzymes [15]. FPP and GGPP are further elongated by long-chain prenyl diphosphate synthases, and their products play important roles in the synthesis of essential molecules such as ubiquinones, menaquinones, some hemes, hormones, and vitamins. Very little is known about the role of the longer-chain isoprenoid biosynthesis enzymes in *T. gondii*.

The number of isoprenyl units that are incorporated into the tail of UQ differ between species [6], for example, *P. falciparum* synthesizes UQ_8_ [38] while humans synthesize UQ_10_, and *T. cruzi* synthesizes UQ_9_ [26]. The synthesis of UQ and its insertion in the mitochondrial membrane is unexplored in *T. gondii* or any other apicomplexan parasite. Considering that the mitochondrion of *T. gondii* and also of other apicomplexan parasites is critical for their replication and several major antiparasitic drugs, such as atovaquone [39] and endochin-like quinolones [40], inhibit the mitochondrial electron transport chain at the coenzyme Q:cytochrome c reductase, we explored TgCoq1 as a potential target for chemotherapy. Endochin-like quinolones target mitochondrial respiration, and they were found to eliminate tissue cysts at ∼76-88% of the control [41]. Bisphosphonates represent a promising class of drug candidates and they have been used previously to treat other human diseases like osteoporosis [11]. Work from our laboratory and others have shown that the isoprenoid pathway is a validated target for controlling *Toxoplasma* [7, 14, 42–46], *Plasmodium spp.*[42, 47–49] and *Cryptosporidium* [50].

Screening of several bisphosphonate derivatives yielded the lipophilic bisphosphonate BPH-1218 that inhibited the activity of TgCoq1 and *in vitro* growth of *T. gondii*. In addition, BPH-1218 protected mice against a lethal infection with *T. gondii*. Targeting of TgCoq1 by BPH-1218 was underscored by several pieces of evidence. First the sensitivity of the recombinant enzyme with an IC_50_ of 60 nM; second, overexpression of the *TgCoq1* gene resulted in parasites more resistant to its inhibition; third, the inhibition of growth was rescued by the UQ derivative UQ_6_, the EC_50_ of BPH-1218 increasing when an ubiquinone was added to the culture. Compared with other compounds which also showed increased EC_50_ in the presence of UQ_6_, BPH-1218 was the only one for which the protection against inhibition by UQ_6_ was large—almost 8 times.

In conclusion, we present a new *T. gondii* target, TgCoq1, that can be inhibited by specific compounds that are neither toxic nor affect host cell growth. Inhibition of TgCoq1 impacted mitochondrial activity, a validated target in all apicomplexans. The ubiquinone synthesis pathway emerges as a novel and unexplored pathway suitable for new chemotherapeutic approaches.

## MATERIALS AND METHODS

### Ethics Statement

All animal care and therapy studies were carried out in accordance with the NIH guidelines. The animal use protocol was reviewed and approved by the Institutional Animal Care and Use Committee (IACUC) of the University of Georgia. AUP# A2018 02-021.

### Phylogenetic Analysis

The protein sequences were retrieved from the National Center for Biotechnology Information (NCBI) database. The phylogenetic tree from Li et al [51] was used as a guide to search for known solanesyl diphosphate synthases (SPPS), hexaprenyl diphosphate synthases (HexPPS), and other long chain prenyl diphosphate synthases. The *T. gondii* Coq1 sequence was used to query the NCBI database using the genomic Basic Local Alignment Search Protein (BLASTP) tool by selecting organisms ranging from prokaryotes to eukaryotes. The default BLAST parameters (expected threshold: 10, low-complexity filter, BLOSSUM 62 substitution matrix) were used. Alignments for phylogenetic trees were constructed using phylogeny.fr website [52, 53]. Protein sequences were inputted to create multiple sequence alignments using the ClustalW program with the auto strategy settings [54]. The alignments were then trimmed manually to trim the extended N-terminals of the TgCoq1, HhPPS, and the SnPPS. The trimmed sequence alignments were then used to create a maximum likelihood tree through Molecular Evolutionary Genetics Analysis (MEGA) with the Jones-Taylor-Thornton (JTT) matrix-based model [55] with 1000 bootstraps and branch lengths measured in the number of substitutions per site [56, 57].

### Chemicals and reagents

Oligonucleotide primers were obtained from Integrated DNA Technologies (IDT). Taq DNA polymerases were from Denville Scientific Inc. and Invitrogen. TRIzol reagent, SuperScript TM III Reverse Transcriptase, TOPO TA cloning kit, and GeneRacer™ Advanced RACE Kit were from Invitrogen and dNTP were from New England BioLab Inc. Restriction enzymes were from New England BioLab Inc. and Promega. Plasmid miniprep kit, Gel extraction kit and DNA purification kit were from Zymo Research. Secondary antibodies with fluorescence for the western blots are from Licor. IPP, DMAPP, GPP, FPP, GGPP were from Sigma. [4-^14^C] Isopentenyl diphosphate triammonimum salt (55.0 mCi/mml) was from PerkinElmer Life Sciences. Ubiquinone Q6 (UQ_6_) and Q8 (UQ_8_) were from Avanti Polar Lipids. Ubiquinone Q10 (UQ_10_) was from Sigma-Aldrich. Silica gels HPTLC were from Analtech. All other reagents were analytical grade or better.

### Cultures

*T. gondii* RH strain tachyzoites were cultured using hTERT (human telomerase reverse transcriptase) cells with 1% BCS and purified as described earlier [58, 59]. Host cells were grown in Dulbecco’s modified minimal essential medium supplemented with 10 % cosmic calf serum. Cell cultures were maintained at 37°C with 5 % CO_2_.

### Protein Studies and Enzymatic Determinations

The TgCoq1 cDNA without the N-terminal 453 nucleotides was cloned into the NcoI and XhoI sites in the pET28a vector. The expression cassette was introduced into *E. coli* BL21-Codon (+) cells. The transformed *E. coli* were selected with Kanamycin. Expression of the recombinant protein was optimally induced by adding 0.4 mM isopropyl β-thiogalactopyranoside (IPTG) to the culture at an OD_600_ of 0.5-0.8. The bacterial culture was grown overnight at 18°C. A HisBind 900 cartridge from Novagen was used to purify the recombinant protein as per the manufacturer’s instructions. The purified protein was desalted using a His Trap Desalting columns from GE Healthcare according to the prescribed protocol and was stored at −80°C with 40 % glycerol.

Kinetic parameters for the recombinant TgCoq1 were calculated from the activity obtained by varying the concentration of the allylic substrates (GPP, FPP, GGPP and IPP) and keeping concentration of counter substrates at a saturating level. The nonlinear regression analysis in Sigma plot 10.0 was used to calculate the Kinetic parameters.

For the enzymatic activity, a standard protocol for medium/long chain prenyl diphosphate synthase was followed as described previously [26]. The enzyme activity was measured by assessing the amount of [4-^14^C] IPP incorporated into butanol-extractable polyprenyl diphosphates. The standard assay mixture contained, in a final volume of 100 µL, Tris-HCl (100 mM, pH 7.4) 100 µM of any of the allylic substrates FPP/GGPP/GPP, 1 mM MgCl_2_,1 mM DTT, 1 % v/v Triton X-100, 100 µM [4-^14^C]-IPP (1 µCi/µMol) and 500 ng of purified protein. Reactions were incubated at 37°C for 30 min unless otherwise indicated. The radioactive prenyl products were extracted with 1-butanol, washed with NaCl-saturated water and activities were calculated from the DPM values.

### Product analysis by reverse phase thin-layer chromatography

The radioactive prenyl diphosphate products were extracted from 500 µL reaction and hydrolyzed to their corresponding alcohols (polyprenols) by using potato acid phosphatase. The enzymatic hydrolysis reaction was set at 37°C overnight as described before [16]. The resultant products were extracted with n-pentane and separated on a reverse-phase TLC plate with acetone:water (19:1 v/v) as the solvent system. The positions of the polyprenol standards were visualized by iodine vapors and marked with pencils. The radioactive polyprenol products were exposed and visualized by autoradiography.

### Extractions and UQ_7_ measurements

Ubiquinone (UQ) from *T. gondii* was isolated using a standard modified protocol as follows, [6]. Parasites were collected and purified by filtration through an 8 μM, 5 μM, and 3 μM nucleopore membrane, followed by two wash steps in Buffer A with glucose (116 mM NaCl, 5.4 mM KCl, 0.8 mM MgSO_4_.7H_2_O, 50 mM HEPES, 5.5 mM Glucose), and resuspended in a hypoosmotic buffer (20 mM HEPES-Tris pH 7.4, 1 mM EDTA) and centrifuged. The final parasite pellet was subject to 4 freeze-thaw cycles of liquid nitrogen for 5 min and 37°C for 1 minute, to lyse cells. The cell pellet was resuspended in 500 μl of a 1-propanol:water (3:1) followed by 2 cycles of 60 s of grinding with glass beads (425-600 μm). The slurry was transferred to a tube and an additional 500 μl of the 1-propanol:water plus 500 μl of n-hexane was added to extract the UQ, vortexed, and centrifuged at 13,000 rpm for 5 minutes. The upper phase containing the UQ was collected into a glass tube and the grinding was repeated. The UQ containing phase was dried under Nitrogen gas and analyzed by TLC and HPLC.

### TLC, HPLC and Mass Spec analysis

The UQ extract was analyzed by normal-phase thin layer chromatography (TLC) with authentic UQ_6_, UQ_8_ and UQ_10_ standards. The samples of UQ extracts were solubilized in chloroform/methanol (2:1, v/v) for spotting. Normal-phase TLC was conducted on a Kieselgel 60 F_254_ plate (Merck Millipore) and was developed with benzene for separation for 1 hour. The plate was viewed under UV illumination, the UQ band was collected, and the sample was extracted with hexane/isopropanol (1:1, v/v). Samples were vacuum-dried at 50°C and solubilized in ethanol. Purified UQ was subjected to high-performance liquid chromatography on a Shimadzu HPLC Class VP series equipped with a reverse phase YMC-Pack ODS-A column (YMC). Ethanol was used as the mobile phase at a flow rate of 1.0 mL/min, and detection was performed by monitoring absorption at 275 nm. Toxoplasma UQ_7_ was pinpointed comparing with UQ_7_ standard directly obtained and purified from *E. coli* harboring heptaprenyl diphosphate synthase gene of *H. influenzae* (*E. coli* KO229/pMN18).

The quantification of UQ_7_ of toxoplasma was done by using 10 µg of UQ_10_ as an internal control. UQ_7_ and UQ_10_ were co-purified with hexane/isopropanol (1:1, v/v). The quantification of UQ_7_ was calculated based on the peak area of UQ_7_ and UQ_10_ in the HPLC chromatogram and cell numbers of parasites.

For LC/MS analysis, toxoplasma UQ_7_ extract was solubilized with methanol/isopropanol (4:1 v/v) with 0.01 µM of ammonium formate and subjected to Waters Xevo TQMS coupled with Acquity UPLC system with Acquity UPLC BEH C18 column. Methanol/isopropanol (4:1 v/v) with 0.01 µM of ammonium formate was used as mobile phase at the flow rate of 0.3 mL/min. The detection was performed by monitoring absorption at 275 nm PDA (photo diode array), and total ion chromatogram (TIC) was extracted for specific UQ_7_ peak (MS scan). The MS spectra showed the specific m/z value of UQ_7_ in the form of ammonium adduction.

### Genetic manipulations

For *in situ* tagging, a fragment of approximately 2 kbs was amplified from the genomic locus (3’ region) of the *TgCoq*1 gene using primers 1 and 2 (Table S5). The fragment was cloned in the pLic-3HA-CAT plasmid [28] and the construct was linearized with *NheI* for transfection of *TatiΔku80* parasites. Clonal cell lines were generated after selection and subcloning and termed *TgCoq1-3HA*. A promoter insertion plasmid was generated by cloning two fragments from the 5’ end of the *TgCoq1* gene into the pDT7S4myc plasmid [28]. One fragment corresponds to the *TgCoq1* 5’ flanking region (predicted promoter/5’UTR) and was amplified with primers 3 and 4 (Table S5, underlined sequences correspond to *NdeI* restriction sites). The second fragment corresponds to the 5’ *TgCoq1* coding sequence beginning with the start codon, which was amplified with primers 5 and 6 (Table S5) (underlined sequences correspond to BglII and AvrII restriction sites). The plasmid was linearized with AvrII for transfection of *TatiΔku80* and *TgCoq1-3HA* cells. The clonal lines created after selection and subcloning were termed *iΔCoq1* and *iΔCoq1-3HA*.

The full-length cDNA of *TgCoq1* was amplified with primers TgCoq1-PI-BGLII-F (primer 6) and TgCoq1-PI-AVRII-R (primer 7). The RT-PCR product was cloned in the Zero blunt Topo vector from Invitrogen. After the sequence was verified, the *TgCoq1* ORF was removed by digestion with BglII and AvrII, cloned into the BglII and AvrII sites of pCTH3 [60] to generate the **pCTCoq1HA** plasmid for complementation. Cosmid **PSBME30** (toxodb.org), which contains the *TgCoq1* genomic locus was obtained from Dr. L. David Sibley (Washington University). For the complementation with the *Trypanosoma cruzi* solanesyl diphosphate synthase (TcSPPS), 453 bp of the 5’ end of the *TgCoq1* ORF was fused with the cDNA of TcSPPS, and the pCTH3 vector by Gibson Assembly to make the p**CTSPPSHA** construct. These three constructs were used for complementation of the conditional knockout cells. The complemented cells were selected by ATc. After surviving ATc selection, the cells were subcloned by limiting dilution. The single clones were analyzed by PCR, and confirmed by Southern blot.

### Immunofluorescence (IFA), western blots, and Southern blots

For IFAs of intracellular parasites subconfluent HFF host cells were used and infected with tachyzoites for 24 hours and fixed with 3% paraformaldehyde. For IFAs of extracellular parasites, *T. gondii* tachyzoites were released by lysing hTERT cells and fixed in 3% paraformaldehyde for 1 h at room temperature. Parasites were adhered to poly-L-Lysine coated cover slips. In both cases parasites were permeabilized with 0.3 % Triton X-100 for 30 min and blocked with blocking buffer (3 % Bovine serum albumin). Incubations with primary and secondary antibodies were for 1 h each. Primary anti-HA antibodies were used at a dilution of 1:200. Secondary antibodies, in all cases, were used at a concentration of 1:1,000. Cover slips with parasites were mounted on glass slides with Antifade-DAPI. A Delta Vision fluorescence microscope was used to observe cells and a Photometrics Coolsnap camera to capture images. Deconvolved images were obtained by using softWoRx deconvolution software.

For western blots, separated proteins in SDS page were transferred to a nitrocellulose membrane and blocked overnight at 4°C with 5 % nonfat milk in PBS-T (0.1 % Tween-20 in PBS). The protein bands were developed by autoradiography on an X-ray film using an ECL detection kit or imaged on a Licor Odyssey CLx machine using secondary antibodies with fluorescence at either 680 or 800 nm. Note that the monoclonal anti-HA antibody used in Fig. 3C is from Covance while the one used in Fig. S4C is a gift from Dr. Chris West.

*T. gondii* genomic DNA was purified and digested with EcoRI or EcoRV for Southern blot analysis. The probe used in Southern blot is the purified PCR product of the 453 bp fragment at the 5’ of TgCoq1 cDNA. The probe was labeled with ^32^P by random priming.

### TgCoq1 antibody production

The TgCoq1 recombinant protein was generated with a *E. coli* expression system and the whole gene was cloned in the pET-28a (+) vector. The *TgCoq1* coding region was released from the pCR-2.1-TOPO vector by digestion with NcoI and NotI and ligated into the NcoI and NotI sites of the pET-28a (+) vector. The resultant pET-28a (+)-TgCoq1 expression construct was modified with a C-terminal 6x His-tag to produce a His-tagged fusion protein. The construct was sequenced and used for transformation of *E. coli* BL21 codon plus. Purified protein (100 μg) was first mixed with Freund’s complete adjuvant and used to inoculate mice which were boosted three times 2 weeks apart with 50 μg peptide in Freund’s incomplete adjuvant [61, 62]. The final serum was tested on *Coq1-HA* lysates and compared with anti-HA to confirm size and antibody purity. The anti-Coq1 serum was affinity purified prior to use for IFAs and westerns. Work with mice was carried out in strict accordance with the Public Health Service Policy on Humane Care and Use of Laboratory Animals and Association for the Assessment and Accreditation of Laboratory Animal Care guidelines. The animal protocol was approved by the University of Georgia’s Committee on the Use and Care of Animals (protocol A2018 02-021). All efforts were made to humanely euthanize the mice after collecting blood.

### Plaque assays and plaquing efficiency

Plaque assays were performed with confluent hTERT cells in six-well plates infected with 150 parasites per well and incubated with or without ATc [62]. After 8 days of incubation, parasites were fixed with 100% ethanol and stained as previously described [63]. ImageJ software was used for quantification of plaque size. Fifteen plaques per well for each biological replicate were counted and averaged. Plaquing efficiency assays were performed with confluent hTERT host cells in six-well plates infected with 2,000 parasites per well. Parasites were preincubated for 3 days with ATc, and then incubated with or without ATc. Thirty minutes after inoculation parasites are washed off with PBS and replaced with fresh DMEM media. After 5 days of incubation the parasites were fixed and stained using the same protocol as the plaque assays. The number of plaques were counted and averaged from 3 biological replicates. For the plaque assays measuring growth rescue we plated 75 parasites per well in a 12-well plate with or without ATc. At 7 days we removed the media, washed, and replaced with media without ATc and allowed the parasites to grow for an additional 7 days. We also included controls without ATc and with ATc for the full 14 days and also changed the media in those wells at day 7. We also tested a plate without washing at day 7 for the with/without ATc and saw the same result as with washing. The same was performed for the BPH-1218 at both 1 µM and 2 µM concentrations. The size of the plaques were measured using Fiji and averaged from 3 biological replicates.

### Measuring mitochondrial membrane potential and oxygen consumption

The mitochondrial membrane potential was measured by the safranine method according to [32] and [35]. Freshly lysed parasites were collected and filtered through an 8 µm filter to remove host cell debris. The parasites were washed twice with BAG and resuspended at 1 x 10^9^ parasites per ml. An aliquot of 50 µl of the parasite suspension (5 x 10^7^ cells) was added to a cuvette containing Safranine O (2.5 µM) and succinate (1 mM) in 2 ml of reaction buffer (125 mM sucrose, 65 mM KCl, 10 mM HEPES-KOH pH 7.2, 1 mM MgCl_2_, and 2.5 mM K_3_PO_4_ pH 7.2). The cuvette was placed in a Hitachi F-7000 fluorescence spectrophotometer and digitonin (30 µM) was added to selectively permeabilize the plasma membrane. The cells were then allowed to equilibrate, and ADP (10 µM final) was added to stimulate oxidative phosphorylation and oligomycin (2 µg/ml final) to inhibit the ATP synthase. FCCP (5 µM) was used to depolarize the mitochondrial membrane. The quantification of the mitochondrial membrane potential is the difference of the maximum fluorescence (after addition of FCCP) minus the minimum fluorescence (before addition of ADP).

Measurements of oxygen consumption are performed with an Oroboros O_2_kFluoRespirometer and the data was analyzed using DatLab software and GraphPad Prism 7 following a published protocol [64]. The parasites were collected using the same protocol described above for mitochondrial membrane potential. 5 x 10^7^ parasites were loaded into the chamber with succinate (5 mM), BSA (0.2%), EGTA (50 µM) in 2 ml of reaction buffer (125 mM Sucrose, 65 mM KCl, 10 mM HEPES-KOH, 2.5 mM potassium phosphate). The chamber was closed to minimize the effects of oxygen from the environment and the suspension was allowed to equilibrate for 10 minutes. Digitonin (12.5 µM) was added and the suspension allowed to equilibrate for 10 min, then ADP (200 µM) was added followed by oligomycin (1.5 µM) after 5-6 min. The quantification in Fig. 5C shows the average peaks after each addition of drug for 3 biological replicates for each cell type. The basal levels represent the normal oxygen consumption of the mitochondria after digitonin. The Ox. Phos. represents the average oxidative phosphorylation after the addition of ADP. The Min. Ox. Phos. represents the minimum oxidative phosphorylation after the addition of oligomycin.

### *In vitro* drug screening and growth assays

To test the effect of bisphosphonates on TgCoq1 activity, enzymatic reactions were set up in the presence of different bisphosphonate compounds. Preliminary screening for potential inhibitors was carried out at a final concentration of 1 µM and 10 µM, respectively and IC_50_s were calculated for the ones that showed > 50% inhibition at 1 µM. The protocol used was previously described [65].

Experiments on *T. gondii* tachyzoites were carried out using parasites expressing red fluorescent protein (RFP) [66] with the modification described by Recher et al [67]. Tachyzoites were maintained in human fibroblasts (hTERT cells). For drug testing and growth assays, parasites were purified by passing them through a 25-gauge needle, followed by filtration through a 3 μm filter. Human fibroblasts were cultured in 96-well black sided plates for 24 hours prior to the addition of 4,000 fluorescent tachyzoites/well. Fluorescence values were measured for up to 9 days, and both excitation (544 nm) and emission (590 nm) were read from the bottom of the plates in a Molecular Devices plate reader [46]. The EC_50_s were calculated using Prism software. The UQ_6_ experiments were performed using the same protocol as described above. The parasites were supplemented with 10 μM UQ_6_. The drug compounds were tested at 2X or 4X their known EC_50_ values.

#### Cytotoxicity to hTert cells

The cytotoxicity was tested using an alamarBlue assay as described by Recher et al. (66). Toxicity was not high enough at the concentrations of drugs tested for detection with alamarBlue.

### *In vivo* drug screening

Experiments were carried out as described previously [49] using 100 *T. gondii* tachyzoites of the RH strain to infect Webster mice. Drugs were dissolved in 10% kolliphor HS 15 and were inoculated intraperitoneally (i.p.). Treatment was initiated 6 hours after infection and administered daily for 10 days.

For the imaging experiments, 20 *T. gondii* tachyzoites of the RH strain (RH-cLuc-GFP, a gift from Saeij lab) were used to infect Webster mice. Treatment protocol was similar to the described above. *In vivo* imaging of infected animals was performed at day 8. The δ-Luciferin potassium salt (Gold Biotech) was dissolved at a concentration of 15.4 mg/ml in sterile PBS. 200 ml of this 8-Luciferin potassium salt solution was injected into each mouse via an intraperitoneal (i.p.) route [34]. The mice were anesthetized using 2.5% (vol/vol) gaseous isofluorane in oxygen. Then the mice were imaged on an IVIS 100 imager (Xenogen, Alameda, CA) [68]. Quantification of bioluminescence result was performed using Living Image v4.3 software (Xenogen).

### Statistical Analysis

Experimental data are expressed as the mean values ± standard deviations (SD) from at least three independent experiments unless indicated otherwise. Statistical analyses were performed by Student’s t-test using GraphPad PRISM version 8.2 for experiments with two experimental conditions. For experiments with 3 or more conditions a two-way anova statistical analysis was used (also stated in the figure legends). The results of the *in vivo* studies were analyzed using the Kaplan-Meier log rank test (GraphPad Prism 7 software). A *P* value of < 0.05 was considered statistically significant.

## ACKNOWLEDGEMENTS

We thank Rodrigo Baptista for advice on the phylogenetic analysis, Mayara Bertolini for training on the use of the Oroboro, Andrea Hortua Triana for assistance with the generation of the anti-TgCoq1 antibodies. Juan Camilo Arenas Garcia helped with the alphafold prediction. The anti-TOM40 antibodies were a gift from Boris Striepen. A monoclonal anti-HA antibody was a gift from Chris West. The work was funded by NIH grants AI102254 and AI147661 to SNJM. MAS was partly funded through a T32 Training Grant fellowship T32AI060546. MK was funded by a grant-in-aid from the Ministry of Education, Culture, Sports, Science, and Technology of Japan [#21H02117]. EO was supported by the University of Illinois Foundation and by a Harriet A. Harlin Professorship.

## AUTHORS CONTRIBUTION

MAS and ZHL performed most experiments, analyzed the data, and wrote the manuscript. RB cloned and characterized the recombinant enzyme. BB performed some of the UQ_6_ drug testing. CL performed the initial characterization of the mutant. JJ and MK analyzed the toxoplasma extracts for the determination of UQ_7_. SRM synthesized all the bisphosphonates and carried out computational structure predictions. EO edited the manuscript. SNJM supervised the project and wrote and edited the manuscript.

## REFERENCES

1. Hill, D. and J.P. Dubey, Toxoplasma gondii: transmission, diagnosis and prevention. Clin Microbiol Infect, 2002. 8(10): p. 634–40.

2. Weiss, L.M. and J.P. Dubey, Toxoplasmosis: A history of clinical observations. Int J Parasitol, 2009. 39(8): p. 895–901.

3. Guo, M., A. Mishra, R.L. Buchanan, J.P. Dubey, D.E. Hill, H.R. Gamble, J.L. Jones and A.K. Pradhan, A Systematic Meta-Analysis of Toxoplasma gondii Prevalence in Food Animals in the United States. Foodborne Pathog Dis, 2016. 13(3): p. 109–18.

4. Alday, P.H. and J.S. Doggett, Drugs in development for toxoplasmosis: advances, challenges, and current status. Drug Des Devel Ther, 2017. 11: p. 273–293.

5. Porter, S.B. and M.A. Sande, Toxoplasmosis of the central nervous system in the acquired immunodeficiency syndrome. N Engl J Med, 1992. 327(23): p. 1643–8.

6. Okada, K., K. Suzuki, Y. Kamiya, X. Zhu, S. Fujisaki, Y. Nishimura, T. Nishino, T. Nakagawa, M. Kawamukai and H. Matsuda, Polyprenyl diphosphate synthase essentially defines the length of the side chain of ubiquinone. Biochim Biophys Acta, 1996. 1302(3): p. 217–23.

7. Nair, S.C., C.F. Brooks, C.D. Goodman, A. Sturm, G.I. McFadden, S. Sundriyal, J.L. Anglin, Y. Song, S.N. Moreno and B. Striepen, Apicoplast isoprenoid precursor synthesis and the molecular basis of fosmidomycin resistance in Toxoplasma gondii. J Exp Med, 2011. 208(7): p. 1547–59.

8. Gershenzon, J. and N. Dudareva, The function of terpene natural products in the natural world. Nat Chem Biol, 2007. 3(7): p. 408–14.

9. Imlay, L. and A.R. Odom, Isoprenoid metabolism in apicomplexan parasites. Curr Clin Microbiol Rep, 2014. 1(3-4): p. 37–50.

10. Oldfield, E., Targeting isoprenoid biosynthesis for drug discovery: bench to bedside. Acc Chem Res, 2010. 43(9): p. 1216–26.

11. Roelofs, A.J., K. Thompson, S. Gordon and M.J. Rogers, Molecular mechanisms of action of bisphosphonates: current status. Clin Cancer Res, 2006. 12(20 Pt 2): p. 6222s–6230s.

12. Johnston, S., K. Wilson, H. Varker, E. Malangone-Monaco, P. Juneau, E. Riehle, S. Satram-Hoang, N. Sommer and S. Ogale, Real-world Direct Health Care Costs for Metastatic Colorectal Cancer Patients Treated With Cetuximab or Bevacizumab-containing Regimens in First-line or First-line Through Second-line Therapy. Clin Colorectal Cancer, 2017. 16(4): p. 386–396 e1.

13. Lange, B.M., T. Rujan, W. Martin and R. Croteau, Isoprenoid biosynthesis: the evolution of two ancient and distinct pathways across genomes. Proc Natl Acad Sci U S A, 2000. 97(24): p. 13172–7.

14. Ling, Y., Z.H. Li, K. Miranda, E. Oldfield and S.N. Moreno, The farnesyl-diphosphate/geranylgeranyl-diphosphate synthase of Toxoplasma gondii is a bifunctional enzyme and a molecular target of bisphosphonates. J Biol Chem, 2007. 282(42): p. 30804–16.

15. Li, Z.H., S. Ramakrishnan, B. Striepen and S.N. Moreno, Toxoplasma gondii relies on both host and parasite isoprenoids and can be rendered sensitive to atorvastatin. PLoS Pathog, 2013. 9(10): p. e1003665.

16. Koyama, T., Molecular analysis of prenyl chain elongating enzymes. Biosci Biotechnol Biochem, 1999. 63(10): p. 1671–6.

17. Liang, P.H., T.P. Ko and A.H. Wang, Structure, mechanism and function of prenyltransferases. Eur J Biochem, 2002. 269(14): p. 3339–54.

18. Saiki, R., A. Nagata, T. Kainou, H. Matsuda and M. Kawamukai, Characterization of solanesyl and decaprenyl diphosphate synthases in mice and humans. FEBS J, 2005. 272(21): p. 5606–22.

19. Kelley, L.A., S. Mezulis, C.M. Yates, M.N. Wass and M.J. Sternberg, The Phyre2 web portal for protein modeling, prediction and analysis. Nat Protoc, 2015. 10(6): p. 845–58.

20. Jumper, J., R. Evans, A. Pritzel, T. Green, M. Figurnov, O. Ronneberger, K. Tunyasuvunakool, R. Bates, A. Zidek, A. Potapenko, et al., Highly accurate protein structure prediction with AlphaFold. Nature, 2021. 596(7873): p. 583–589.

21. Chen, C.-C.M., Satish R.; Han, Xu; Liu, Weidong; Ma, Lixin; Zhai, Chao; Dai, Longhai; Huang, Jian-Wen; Shillo, Alli; Desai, Janish; Ma, Xianqiang; Zhang, Yonghui; Guo, Rey-Ting and Oldfield, Eric Terpene Cyclases and Prenyltransferases: Structures and Mechanisms of Action. ACS Catalysis 2021. 11(1): p. 290–303.

22. Yokoyama, T., M. Mizuguchi, A. Ostermann, K. Kusaka, N. Niimura, T.E. Schrader and I. Tanaka, Protonation State and Hydration of Bisphosphonate Bound to Farnesyl Pyrophosphate Synthase. J Med Chem, 2015. 58(18): p. 7549–56.

23. Sasaki, D., M. Fujihashi, N. Okuyama, Y. Kobayashi, M. Noike, T. Koyama and K. Miki, Crystal structure of heterodimeric hexaprenyl diphosphate synthase from Micrococcus luteus B-P 26 reveals that the small subunit is directly involved in the product chain length regulation. J Biol Chem, 2011. 286(5): p. 3729–40.

24. Desai, J., Y.L. Liu, H. Wei, W. Liu, T.P. Ko, R.T. Guo and E. Oldfield, Structure, Function, and Inhibition of Staphylococcus aureus Heptaprenyl Diphosphate Synthase. ChemMedChem, 2016. 11(17): p. 1915–23.

25. Gautier, R., D. Douguet, B. Antonny and G. Drin, HELIQUEST: a web server to screen sequences with specific alpha-helical properties. Bioinformatics, 2008. 24(18): p. 2101–2.

26. Ferella, M., A. Montalvetti, P. Rohloff, K. Miranda, J. Fang, S. Reina, M. Kawamukai, J. Bua, D. Nilsson, C. Pravia, et al., A solanesyl-diphosphate synthase localizes in glycosomes of Trypanosoma cruzi. J Biol Chem, 2006. 281(51): p. 39339–48.

27. Saiki, R., A. Nagata, N. Uchida, T. Kainou, H. Matsuda and M. Kawamukai, Fission yeast decaprenyl diphosphate synthase consists of Dps1 and the newly characterized Dlp1 protein in a novel heterotetrameric structure. Eur J Biochem, 2003. 270(20): p. 4113–21.

28. Sheiner, L., J.L. Demerly, N. Poulsen, W.L. Beatty, O. Lucas, M.S. Behnke, M.W. White and B. Striepen, A systematic screen to discover and analyze apicoplast proteins identifies a conserved and essential protein import factor. PLoS Pathog, 2011. 7(12): p. e1002392.

29. Fox, B.A., J.G. Ristuccia, J.P. Gigley and D.J. Bzik, Efficient gene replacements in Toxoplasma gondii strains deficient for nonhomologous end joining. Eukaryot Cell, 2009. 8(4): p. 520–9.

30. Mazumdar, J., H.W. E, K. Masek, A.H. C and B. Striepen, Apicoplast fatty acid synthesis is essential for organelle biogenesis and parasite survival in Toxoplasma gondii. Proc Natl Acad Sci U S A, 2006. 103(35): p. 13192–7.

31. van Dooren, G.G., L.M. Yeoh, B. Striepen and G.I. McFadden, The Import of Proteins into the Mitochondrion of Toxoplasma gondii. J Biol Chem, 2016. 291(37): p. 19335–50.

32. Vercesi, A.E., C.O. Rodrigues, S.A. Uyemura, L. Zhong and S.N. Moreno, Respiration and oxidative phosphorylation in the apicomplexan parasite Toxoplasma gondii. J Biol Chem, 1998. 273(47): p. 31040–7.

33. Shang, N., Q. Li, T.P. Ko, H.C. Chan, J. Li, Y. Zheng, C.H. Huang, F. Ren, C.C. Chen, Z. Zhu, et al., Squalene synthase as a target for Chagas disease therapeutics. PLoS Pathog, 2014. 10(5): p. e1004114.

34. Saeij, J.P., J.P. Boyle, M.E. Grigg, G. Arrizabalaga and J.C. Boothroyd, Bioluminescence imaging of Toxoplasma gondii infection in living mice reveals dramatic differences between strains. Infect Immun, 2005. 73(2): p. 695–702.

35. McPhillie, M.J., Y. Zhou, M.R. Hickman, J.A. Gordon, C.R. Weber, Q. Li, P.J. Lee, K. Amporndanai, R.M. Johnson, H. Darby, et al., Potent Tetrahydroquinolone Eliminates Apicomplexan Parasites. Front Cell Infect Microbiol, 2020. 10: p. 203.

36. Harb, O.S. and D.S. Roos, ToxoDB: Functional Genomics Resource for Toxoplasma and Related Organisms. Methods Mol Biol, 2020. 2071: p. 27–47.

37. Kawamukai, M., Biosynthesis of coenzyme Q in eukaryotes. Biosci Biotechnol Biochem, 2016. 80(1): p. 23–33.

38. de Macedo, C.S., M.L. Uhrig, E.A. Kimura and A.M. Katzin, Characterization of the isoprenoid chain of coenzyme Q in Plasmodium falciparum. FEMS Microbiol Lett, 2002. 207(1): p. 13–20.

39. Srivastava, I.K., H. Rottenberg and A.B. Vaidya, Atovaquone, a broad spectrum antiparasitic drug, collapses mitochondrial membrane potential in a malarial parasite. J Biol Chem, 1997. 272(7): p. 3961–6.

40. Winter, R., J.X. Kelly, M.J. Smilkstein, D. Hinrichs, D.R. Koop and M.K. Riscoe, Optimization of endochin-like quinolones for antimalarial activity. Exp Parasitol, 2011. 127(2): p. 545–51.

41. Doggett, J.S., A. Nilsen, I. Forquer, K.W. Wegmann, L. Jones-Brando, R.H. Yolken, C. Bordon, S.A. Charman, K. Katneni, T. Schultz, et al., Endochin-like quinolones are highly efficacious against acute and latent experimental toxoplasmosis. Proc Natl Acad Sci U S A, 2012. 109(39): p. 15936–41.

42. Martin, M.B., J.S. Grimley, J.C. Lewis, H.T. Heath, 3rd, B.N. Bailey, H. Kendrick, V. Yardley, A. Caldera, R. Lira, J.A. Urbina, et al., Bisphosphonates inhibit the growth of Trypanosoma brucei, Trypanosoma cruzi, Leishmania donovani, Toxoplasma gondii, and Plasmodium falciparum: a potential route to chemotherapy. J Med Chem, 2001. 44(6): p. 909–16.

43. Yardley, V., A.A. Khan, M.B. Martin, T.R. Slifer, F.G. Araujo, S.N. Moreno, R. Docampo, S.L. Croft and E. Oldfield, In vivo activities of farnesyl pyrophosphate synthase inhibitors against Leishmania donovani and Toxoplasma gondii. Antimicrob Agents Chemother, 2002. 46(3): p. 929–31.

44. Szajnman, S.H., G.E. Garcia Linares, Z.H. Li, C. Jiang, M. Galizzi, E.J. Bontempi, M. Ferella, S.N. Moreno, R. Docampo and J.B. Rodriguez, Synthesis and biological evaluation of 2-alkylaminoethyl-1,1-bisphosphonic acids against Trypanosoma cruzi and Toxoplasma gondii targeting farnesyl diphosphate synthase. Bioorg Med Chem, 2008. 16(6): p. 3283–90.

45. Rosso, V.S., S.H. Szajnman, L. Malayil, M. Galizzi, S.N. Moreno, R. Docampo and J.B. Rodriguez, Synthesis and biological evaluation of new 2-alkylaminoethyl-1,1-bisphosphonic acids against Trypanosoma cruzi and Toxoplasma gondii targeting farnesyl diphosphate synthase. Bioorg Med Chem, 2011. 19(7): p. 2211–7.

46. Li, Z.H., C. Li, S.H. Szajnman, J.B. Rodriguez and S.N.J. Moreno, Synergistic Activity between Statins and Bisphosphonates against Acute Experimental Toxoplasmosis. Antimicrob Agents Chemother, 2017. 61(8): p. e02628–16.

47. Singh, A.P., Y. Zhang, J.H. No, R. Docampo, V. Nussenzweig and E. Oldfield, Lipophilic bisphosphonates are potent inhibitors of Plasmodium liver-stage growth. Antimicrob Agents Chemother, 2010. 54(7): p. 2987–93.

48. Jordao, F.M., A.Y. Saito, D.C. Miguel, V. de Jesus Peres, E.A. Kimura and A.M. Katzin, In vitro and in vivo antiplasmodial activities of risedronate and its interference with protein prenylation in Plasmodium falciparum. Antimicrob Agents Chemother, 2011. 55(5): p. 2026–31.

49. Szajnman, S.H., T. Galaka, Z.H. Li, C. Li, N.M. Howell, M.N. Chao, B. Striepen, V. Muralidharan, S.N. Moreno and J.B. Rodriguez, In Vitro and In Vivo Activities of Sulfur-Containing Linear Bisphosphonates against Apicomplexan Parasites. Antimicrob Agents Chemother, 2017. 61(2).

50. Moreno, B., B.N. Bailey, S. Luo, M.B. Martin, M. Kuhlenschmidt, S.N. Moreno, R. Docampo and E. Oldfield, (31)P NMR of apicomplexans and the effects of risedronate on Cryptosporidium parvum growth. Biochem Biophys Res Commun, 2001. 284(3): p. 632–7.

51. Li, Z.H., R. Cintron, N.A. Koon and S.N. Moreno, The N-terminus and the chain-length determination domain play a role in the length of the isoprenoid product of the bifunctional Toxoplasma gondii farnesyl diphosphate synthase. Biochemistry, 2012. 51(38): p. 7533–40.

52. Dereeper, A., S. Audic, J.M. Claverie and G. Blanc, BLAST-EXPLORER helps you building datasets for phylogenetic analysis. BMC Evol Biol, 2010. 10: p. 8.

53. Dereeper, A., V. Guignon, G. Blanc, S. Audic, S. Buffet, F. Chevenet, J.F. Dufayard, S. Guindon, V. Lefort, M. Lescot, et al., Phylogeny.fr: robust phylogenetic analysis for the non-specialist. Nucleic Acids Res, 2008. 36(Web Server issue): p. W465–9.

54. Thompson, J.D., D.G. Higgins and T.J. Gibson, CLUSTAL W: improving the sensitivity of progressive multiple sequence alignment through sequence weighting, position-specific gap penalties and weight matrix choice. Nucleic Acids Res, 1994. 22(22): p. 4673–80.

55. Jones, D.T., W.R. Taylor and J.M. Thornton, The rapid generation of mutation data matrices from protein sequences. Comput Appl Biosci, 1992. 8(3): p. 275–82.

56. Kumar, S., M. Nei, J. Dudley and K. Tamura, MEGA: a biologist-centric software for evolutionary analysis of DNA and protein sequences. Brief Bioinform, 2008. 9(4): p. 299–306.

57. Kumar, S., G. Stecher, M. Li, C. Knyaz and K. Tamura, MEGA X: Molecular Evolutionary Genetics Analysis across Computing Platforms. Mol Biol Evol, 2018. 35(6): p. 1547–1549.

58. Moreno, S.N. and L. Zhong, Acidocalcisomes in Toxoplasma gondii tachyzoites. Biochem J, 1996. 313 ( Pt 2): p. 655–9.

59. Vella, S.A., A. Calixto, B. Asady, Z.H. Li and S.N.J. Moreno, Genetic Indicators for Calcium Signaling Studies in Toxoplasma gondii. Methods Mol Biol, 2020. 2071: p. 187–207.

60. Miranda, K., D.A. Pace, R. Cintron, J.C. Rodrigues, J. Fang, A. Smith, P. Rohloff, E. Coelho, F. de Haas, W. de Souza, et al., Characterization of a novel organelle in Toxoplasma gondii with similar composition and function to the plant vacuole. Mol Microbiol, 2010. 76(6): p. 1358–75.

61. Chasen, N.M., B. Asady, L. Lemgruber, R.C. Vommaro, J.C. Kissinger, I. Coppens and S.N.J. Moreno, A Glycosylphosphatidylinositol-Anchored Carbonic Anhydrase-Related Protein of Toxoplasma gondii Is Important for Rhoptry Biogenesis and Virulence. mSphere, 2017. 2(3).

62. Stasic, A.J., N.M. Chasen, E.J. Dykes, S.A. Vella, B. Asady, V.J. Starai and S.N.J. Moreno, The Toxoplasma Vacuolar H(+)-ATPase Regulates Intracellular pH and Impacts the Maturation of Essential Secretory Proteins. Cell Rep, 2019. 27(7): p. 2132–2146 e7.

63. Roos, D.S., R.G. Donald, N.S. Morrissette and A.L. Moulton, Molecular tools for genetic dissection of the protozoan parasite Toxoplasma gondii. Methods Cell Biol, 1994. 45: p. 27–63.

64. Bertolini, M.S., M.A. Chiurillo, N. Lander, A.E. Vercesi and R. Docampo, MICU1 and MICU2 Play an Essential Role in Mitochondrial Ca(2+) Uptake, Growth, and Infectivity of the Human Pathogen Trypanosoma cruzi. mBio, 2019. 10(3).

65. Ling, Y., G. Sahota, S. Odeh, J.M. Chan, F.G. Araujo, S.N. Moreno and E. Oldfield, Bisphosphonate inhibitors of Toxoplasma gondi growth: in vitro, QSAR, and in vivo investigations. J Med Chem, 2005. 48(9): p. 3130–40.

66. van Dooren, G.G., C. Tomova, S. Agrawal, B.M. Humbel and B. Striepen, Toxoplasma gondii Tic20 is essential for apicoplast protein import. Proc Natl Acad Sci U S A, 2008. 105(36): p. 13574–9.

67. Recher, M., A.P. Barboza, Z.H. Li, M. Galizzi, M. Ferrer-Casal, S.H. Szajnman, R. Docampo, S.N. Moreno and J.B. Rodriguez, Design, synthesis and biological evaluation of sulfur-containing 1,1-bisphosphonic acids as antiparasitic agents. Eur J Med Chem, 2013. 60: p. 431–40.

68. Canavaci, A.M., J.M. Bustamante, A.M. Padilla, C.M. Perez Brandan, L.J. Simpson, D. Xu, C.L. Boehlke and R.L. Tarleton, In vitro and in vivo high-throughput assays for the testing of anti-Trypanosoma cruzi compounds. PLoS Negl Trop Dis, 2010. 4(7): p. e740.

