## Supplemental material for "The heptaprenyl diphosphate synthase (Coq1) is the target of a lipophilic bisphosphonate that protects mice against *Toxoplasma gondii* infection"

**Sleda, Li et al**

**Supplementary Tables and Figures**

**Supplementary Table S1:** Sequences used for the phylogenetic analysis presented in Figure 1.

| Organism_Enzyme | GenBank/EuPathDB ID | Reference |
|---|---|---|
| *Arabidopsis thaliana*_SPPS | NP_177972.2 | (1) |
| *Saccharomyces cerevisiae*_HexPPS | P18900.1 | (2) |
| *Trypanosoma cruzi*_SPPS | EAN82722.1/ Tc00.1047053427091.10 | (3) |
| *Leishmania major*_SPPS | XP_001682016.1/LmjF.15.1020 | Predicted |
| *Plasmodium vivax*_OPPS | VUZ93874.1/PVX_003575 | Predicted |
| *Plasmodium falciparum*_OPPS | XP_001349541.1/PFB0130w | (4) |
| *Toxoplasma gondii*_LONG | XP_018636584.1/TGME49_069430 | This work |
| *Neospora caninum*_PPS | XP_003883948.1/NCLIV_036980 | Predicted |
| *Oryza sativa*_SPPS | Q653T6.1 | (5) |
| *Babesia bovis*_PPS | EDO07087.1 | Predicted |
| *Haemophilus influenzae*_HepPPS | CBY85984.1 | Predicted |
| *Bacillus subtilis* HepPPS | ARW31988.1 | (6) |
| *Listeria monocytogenes* HepPPS | QGK56962.1 | Predicted |
| *Aquifex aeolicus* OPPS | NP_213606.1 | Predicted |
| *Aspergillus fumigatus* HexPPS | EDP53123.1 | Predicted |
| *Mus musculus* SPPS | BAE48219.1 | (7) |
| *Equus caballus* DPPS | XP023487828.1 | Predicted |
| *Homo sapiens* DPPS | BAE48216.1 | (7) |
| *Hammondia hammondi* PPS | HHA_269430/HHA_269430 | Predicted |
| *Cyclospora cayetanensis* PPS | LOC34620908 | Predicted |
| *Sarcocystis neurona* PPS | SN3_03700020 | Predicted |

References


1. Jun L, Saiki R, Tatsumi K, Nakagawa T, Kawamukai M. 2004. Identification and subcellular localization of two solanesyl diphosphate synthases from Arabidopsis thaliana. Plant Cell Physiol 45:1882-8.
2. Ashby MN, Edwards PA. 1990. Elucidation of the deficiency in two yeast coenzyme Q mutants. Characterization of the structural gene encoding hexaprenyl pyrophosphate synthetase. J Biol Chem 265:13157-64.
3. Ferella M, Montalvetti A, Rohloff P, Miranda K, Fang J, Reina S, Kawamukai M, Bua J, Nilsson D, Pravia C, Katzin A, Cassera MB, Aslund L, Andersson B, Docampo R, Bontempi EJ. 2006. A solanesyl-diphosphate synthase localizes in glycosomes of Trypanosoma cruzi. J Biol Chem 281:39339-48.
4. Tonhosolo R, D'Alexandri FL, Genta FA, Wunderlich G, Gozzo FC, Eberlin MN, Peres VJ, Kimura EA, Katzin AM. 2005. Identification, molecular cloning and functional characterization of an octaprenyl pyrophosphate synthase in intra-erythrocytic stages of Plasmodium falciparum. Biochem J 392:117-26.

5. Ohara K, Sasaki K, Yazaki K. 2010. Two solanesyl diphosphate synthases with different subcellular localizations and their respective physiological roles in Oryza sativa. J Exp Bot 61:2683-92.
6. Zhang YW, Koyama T, Marecak DM, Prestwich GD, Maki Y, Ogura K. 1998. Two subunits of heptaprenyl diphosphate synthase of Bacillus subtilis form a catalytically active complex. Biochemistry 37:13411-20.
7. Saiki R, Nagata A, Kainou T, Matsuda H, Kawamukai M. 2005. Characterization of solanesyl and decaprenyl diphosphate synthases in mice and humans. FEBS J 272:5606-22.

**Supplementary Table S2:** physicochemical properties of the TgCoq1 N terminal domain

A

| ${}_{32}$FFSSLSCVSVPPPLCAVS${}_{49}$ | | |
|---|---|---|
| **Physico-chemical properties** | **Polar residues + GLY** | **Nonpolar residues** |
| Hydrophobicity <H> | Polar residues + GLY (n / %) | Nonpolar residues (n / %) |
| 0.888 | 5 / 27.78 | 13 / 72.22 |
| Hydrophobic moment <µH> | Uncharged residues + GLY | Aromatic residues |
| 0.073 | SER 5, GLY 0 | PHE 2 |
| Net charge z | Charged residues | Special residues |
| 0 | | CYS 2, PRO 3 |
| **Hydrophobic face:** C V F P L A | | |

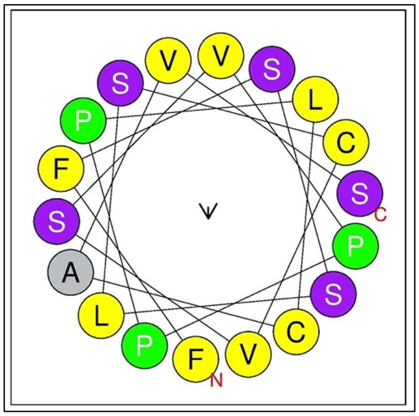

B

| ${}_{68}$VRGEDRRRLSGRRRQVEK${}_{85}$ | | |
|---|---|---|
| **Physico-chemical properties** | **Polar residues + GLY** | **Nonpolar residues** |
| Hydrophobicity <H> | Polar residues + GLY (n / %) | Nonpolar residues (n / %) |
| -0.346 | 15 / 83.33 | 3 / 16.67 |
| Hydrophobic moment <µH> | Uncharged residues + GLY | Aromatic residues |
| 0.212 | GLN 1, SER 1, GLY 2 | |
| Net charge z | Charged residues | Special residues |
| 5 | LYS 1, ARG 7, GLU 2, ASP 1 | CYS 0, PRO 0 |
| **Hydrophobic face:** none | | |

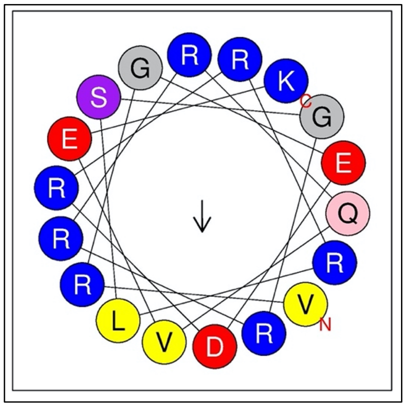

C

| ${}_{185}$EGSEDEGTTSEQPSGGRD${}_{202}$ | | |
|---|---|---|
| **Physico-chemical properties** | **Polar residues + GLY** | **Nonpolar residues** |
| Hydrophobicity <H> | Polar residues + GLY (n /%) | Nonpolar residues (n /%) |
| -0.234 | 17 / 94.44 | 1 / 5.56 |
| Hydrophobic moment <µH> | Uncharged residues + GLY | Aromatic residues |
| 0.082 | GLN 1, SER 3, THR 2, GLY 4 | |
| Net charge z | Charged residues | Special residues |
| -5 | ARG 1, GLU 4, ASP 2, | CYS 0, PRO 1 |
| **Hydrophobic face:** none | | |

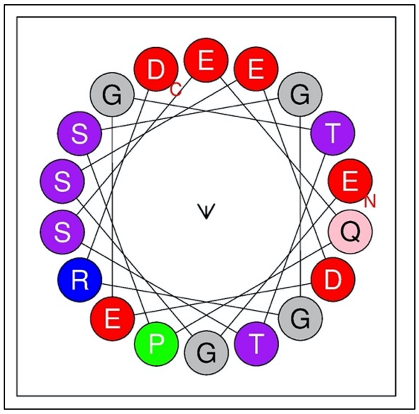

**Supplementary Table S3:** Cytotoxicity of compounds on hTERT cells at 4X and 10X their $EC_{50}$. Values are means ± s.d. of n = 2.

| Compound (µM) | % Inhibition |
|---|---|
| BPH-1218 (4) | 4.30 ± 2.31 |
| BPH-1218 (10) | 10.28 ± 1.07 |
| BPH-1217 (28.4) | 8.01 ± 1.82 |
| BPH-1217 (71) | 6.19 ± 9.55 |
| BPH-1219 (46) | 3.20 ± 2.89 |
| BPH-1219 (115) | 8.01 ± 1.82 |
| BPH-1236 (2.36) | 4.13 ± 1.20 |
| BPH-1236 (5.9) | 9.10 ± 2.38 |
| BPH-1238 (2.72) | 9.77 ± 1.18 |
| BPH-1238 (6.8) | 13.37 ± 6.08 |
| AV (0.12) | 2.33 ± 0.99 |
| AV (0.32) | 2.91 ± 4.90 |
| JAG-21 (0.48) | 6.40 ± 5.50 |
| JAG-21 (1.2) | 5.75 ± 4.98 |

**Supplementary Table S4:** Compounds for CoQ6 rescue experiment from Fig. 7D.

| Compound | Structure | Compound | Structure |
|---|---|---|---|
| **BPH-1218** | 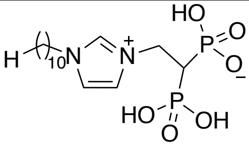 | **BPH-754** | 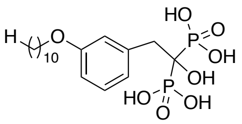 |
| **BPH-1327** | 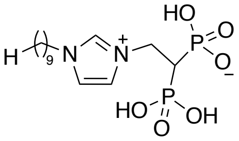 | **Risedronate** | 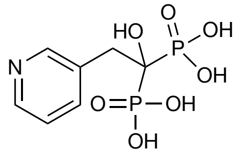 |
| **BPH-1236** | 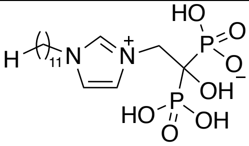 | **SSC-36** | 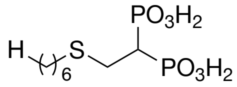 |
| **JAG-21** | 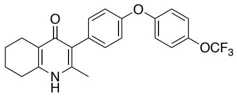 | **CE-22** | 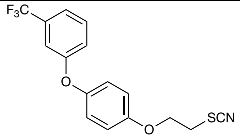 |
| **BPH-1238** | 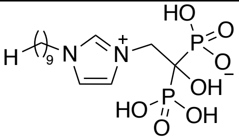 | **MNC-98** | 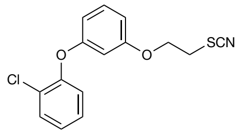 |
| **Atovaquone** | 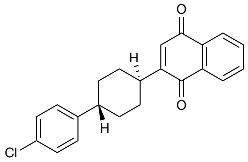 | **CE-29** | 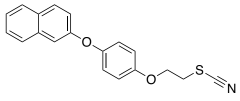 |
| **Pyrimethamine** | 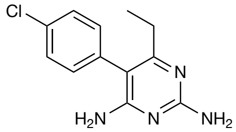 | **MNCA181** | 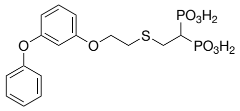 |
| **BPH-1217** | 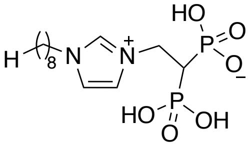 | **CE-109** | 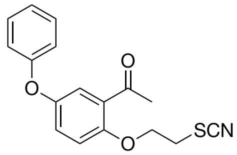 |
| **BPH-1219** | 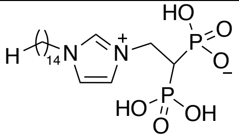 | **CE-91** | 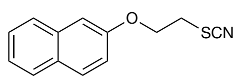 |

**Supplementary Table S5:** Primers used in this work

| Primer | Primer Use | Sequence |
|---|---|---|
| **1** | In situ tagging | tacttccaatccaatttaatgcAGACAACAGCGCAGTCCAAGTCTTG |
| **2** | In situ tagging | tcctccacttccaattttagcCCCAGACCGCCGCTGGAGAGTGGCC |
| **3** | Promoter Insertion 5' flanking sequence fragment | CATATGAAATTAGACAGAAGTGCCGAGAAG |
| **4** | Promoter Insertion 5' flanking sequence fragment | CATATGTTCGCCGACAGACACAAGAGAGATC |
| **5** | Promoter Insertion 5' coding sequence fragment | AGATCTATGACGCTCGTTACTCGACACC |
| **6** | Promoter Insertion 5' coding sequence fragment | CCTAGGGATTCAGAAACGAATTACAGAG |
| **7** | Full-length cDNA F | AGATCTATGACGCTCGTTACTCGACACC |
| **8** | Full-length cDNA R | CCTAGGGATTCAGAAACGAATTACAGAG |

### Supplementary Figures

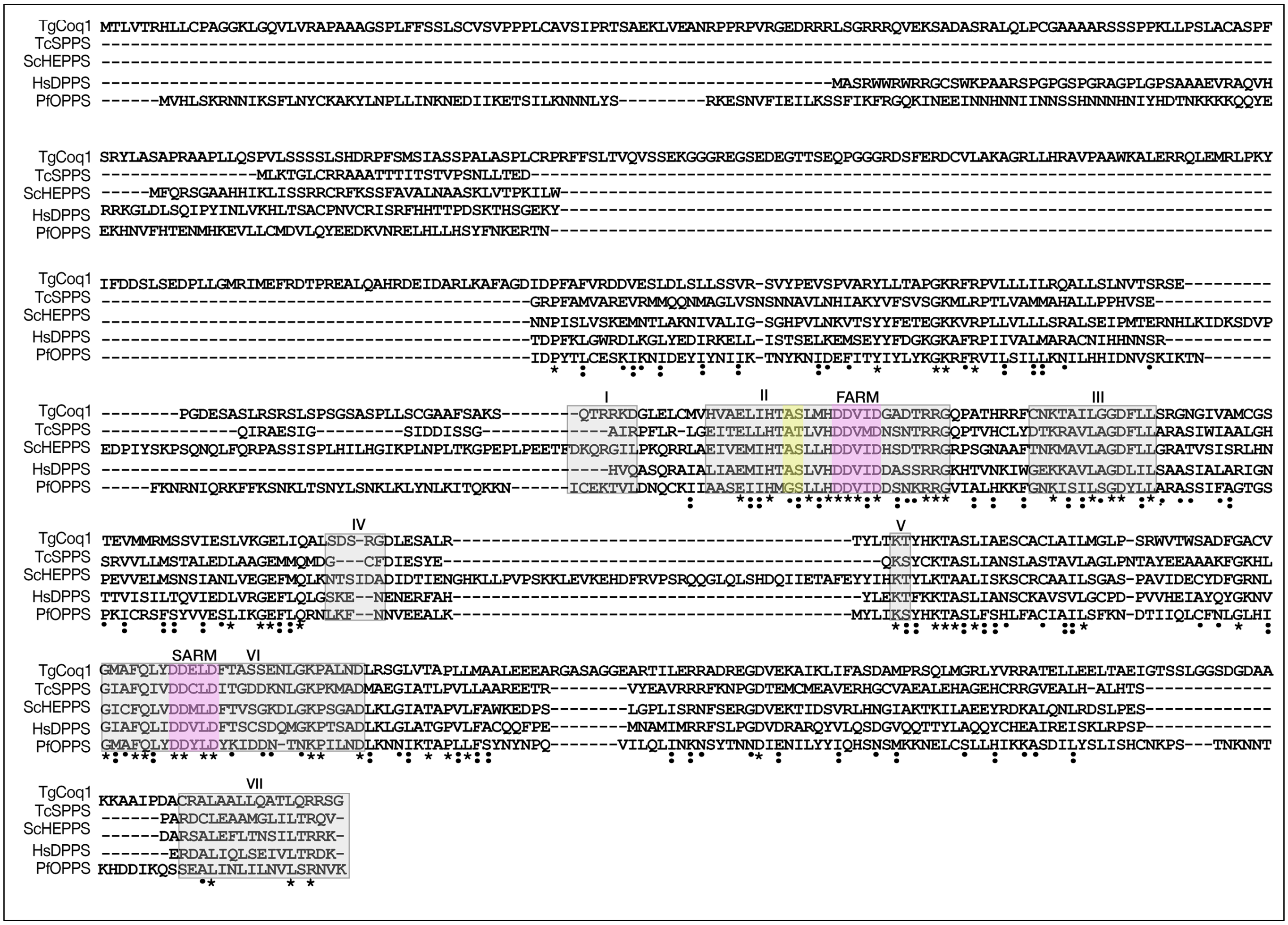


**Supplementary Figure S1.** Clustal sequence alignment of the protein sequences from *T. gondii* Coq1 (TgCoq1), a solanesyl diphosphate synthase from *T. cruzi* (TcSPPS), an octaprenyl diphosphate synthase from *P. falciparum* (PfOPPS), a Hexaprenyl diphosphate synthase from *S. cerevisiae* (SceHexPPS) and the human decaprenyl diphosphate synthase (HsDPPS). The first aspartic rich domain (FARM) and the second aspartic rich domain (SARM) are highlighted in pink and are conserved amongst the sequences. The 4$^{th}$ and 5$^{th}$ positions prior to the FARM are highlighted in yellow. *: highlights identical amino acids between the sequences. The other symbols indicate partial conservation between sequences.

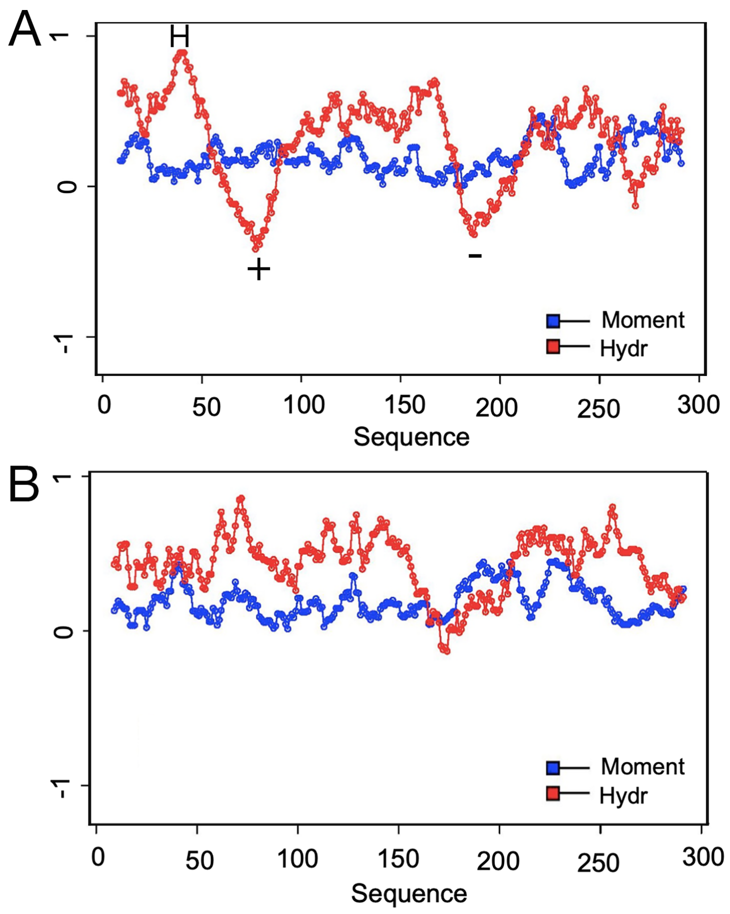


**Supplementary Figure S2. A**, Hydrophobicity <**H**> (red) and hydrophobic moment <**μH**> (blue) as a function of sequence position for the first 300 residues in TgCoq1; H=hydrophobic, the "+" and "-" values indicate the Arg/Lys and Asp/Glu-rich regions. **B**, Hydrophobicity <**H**> (red) and hydrophobic moment <**μH**> (blue) as a function of sequence position for the first 300 residues in TgFPPS.

A

| Trial | $UQ_7$ (µg/1x$10^{10}$parasites) |
|---|---|
| 1 | 4.956 |
| 2 | 3.389 |
| Average | 4.173 |

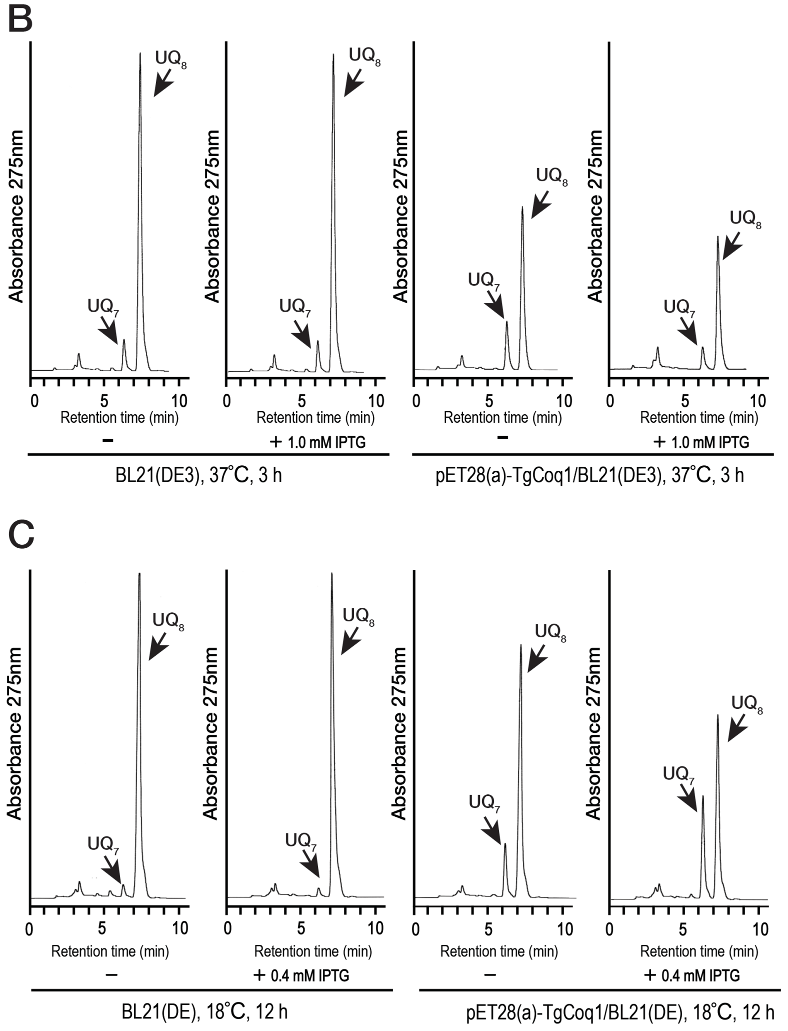


**Supplementary Figure S3.** (A) Amount of Ubiquinone $Q_7$ present in RH wild-type parasites. UQ was measured using HPLC, and the amount of $UQ_7$ present was calculated based on an internal $UQ_{10}$ standard. (B-C) Expression of the entire length of the *TgCoq1* gene cloned in the pET28(a) plasmid under the control of the T7 lac promoter. UQ was extracted from *E. coli* BL21(DE3) harboring pET28(a)-TgCoq1, in which TgCoq1 was induced by addition of isopropyl β-D-thiogalactopyranoside (IPTG) (1.0 mM) at 37°C for 3 hours (B) or 0.4 mM at 18°C for 12 hours (C). They were first separated by TLC and further analyzed by HPLC. These results showed that *E. coli* BL21(DE3) expressing TgCoq1 induced by 0.4 mM IPTG at 18°C for 12 hours clearly produced $UQ_7$ but did not so much by 1.0 mM IPTG at 37 °C for 3 hours.

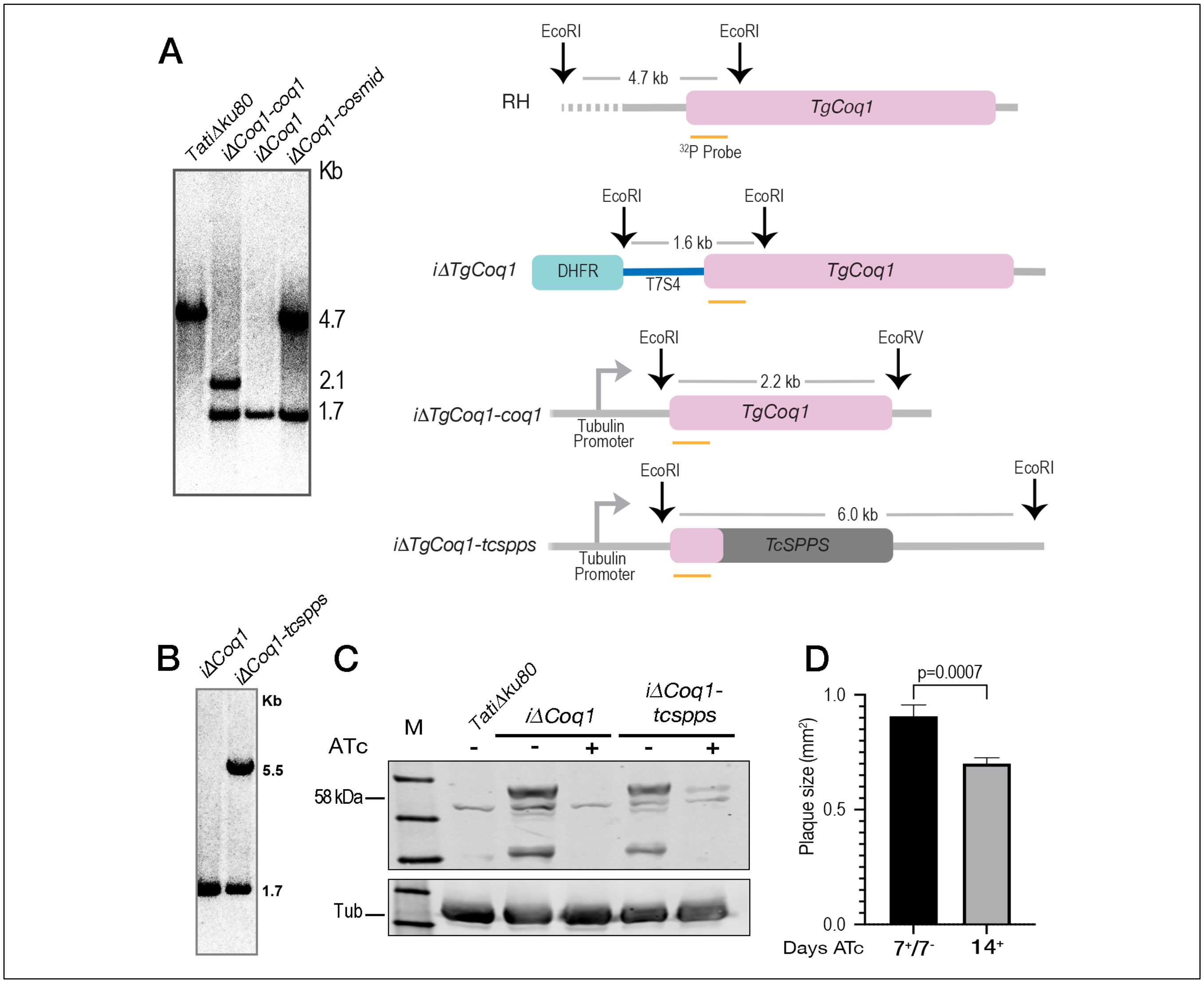


**Supplementary Figure S4.** **A,** Southern blot analysis of genomic DNA isolated from the *iΔCoq1*, *iΔCoq1*-Coq1, and *iΔCoq1*-cosmid complementation mutants. The model depicts the constructs of each of the cell lines, and the location of the enzymes cut sites for the southern blot analysis. **B,** Southern blot analysis of the *iΔCoq1-tcspps* complementation mutant. **C,** Western blot analysis with HA antibody (monoclonal antibody generated at UGA, a gift from Christopher West) showing the HA expression in the *iΔCoq1*-ATc (58 kDA) and the loss of HA signal in the *iΔCoq1* with 3 days ATc. The western also shows the HA signal in the *iΔCoq1-tcspps* (56 kDA). **D,** Plaque size quantification of growth for the *iΔCoq1* cell line grown with ATc for 7 days and then 7 days without ATc or grown for 14 days with ATc. The *iΔCoq1* -ATc cell line was used as a control but was unquantifiable because it lysed the monolayer. (Average from 3 biological replicates, Student's T-test statistical analysis).

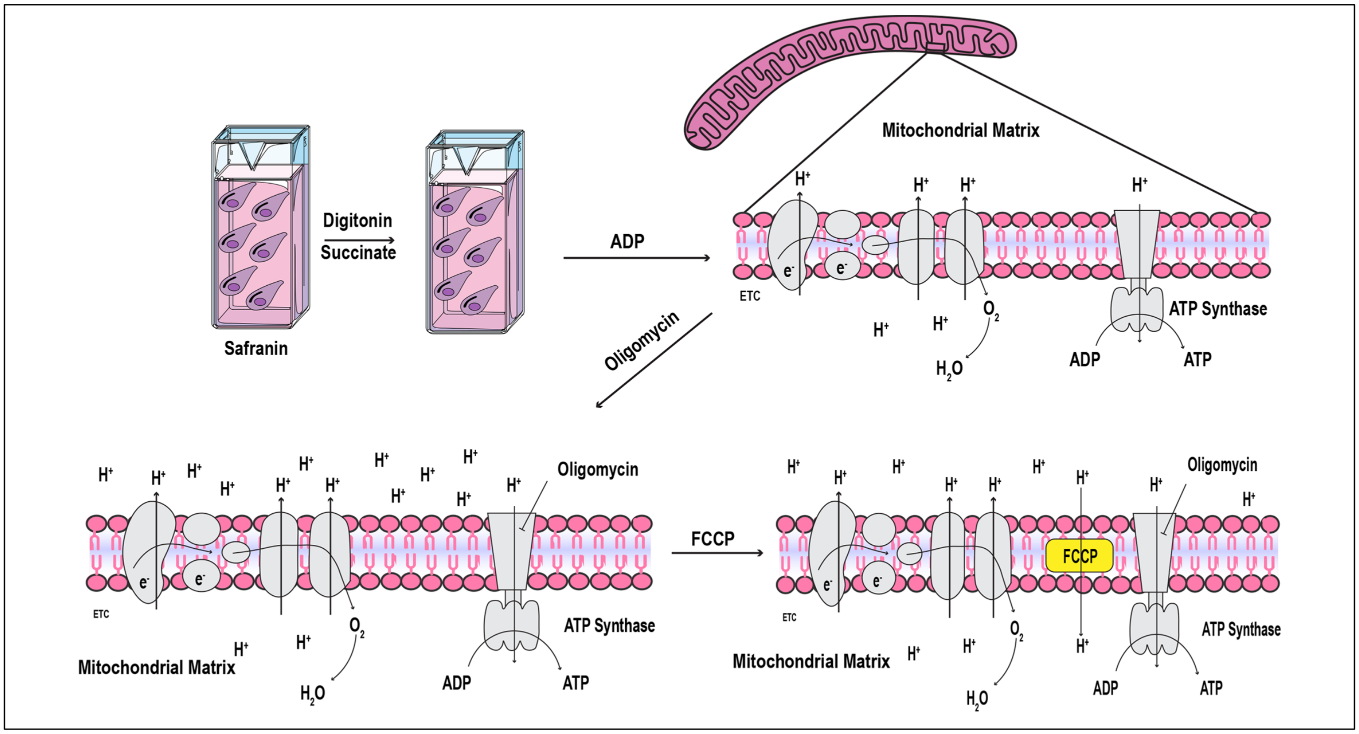


**Supplementary Figure S5.** Model of mitochondrial membrane potential experiments. Digitonin (Dig) addition permeabilizes the plasma membrane for the mitochondrial substrate, succinate. Adenosine diphosphate (ADP) results in synthesis of ATP and use of the proton gradient. Oligomycin (Oligo) inhibits the ATP synthase and allows the membrane potential to recover. Carbonyl cyanide-4-(trifluoromethoxy) phenylhydrazone (FCCP) is a proton ionophore and collapses the $H^+$ gradient.
